# A neural m^6^A/YTHDF pathway is required for learning and memory in *Drosophila*

**DOI:** 10.1101/2020.03.07.982090

**Authors:** Lijuan Kan, Stanislav Ott, Brian Joseph, Eun Sil Park, Crystal Dai, Ralph Kleiner, Adam Claridge-Chang, Eric C. Lai

## Abstract

The roles of epitranscriptomic modifications in mRNA regulation have recently received substantial attention, with appreciation growing for their phenotypically selective impacts within the animal. We adopted *Drosophila melanogaster* as a model system to study m^6^A, the most abundant internal modification of mRNA. Here, we report proteomic and functional analyses of fly m^6^A-binding proteins, confirming nuclear (YTHDC) and cytoplasmic (YTHDF) YTH domain proteins as the major m^6^A binders. Since all core m^6^A pathway mutants are viable, we assessed *in vivo* requirements of the m^6^A pathway in cognitive processes. Assays of short term memory revealed an age-dependent requirement of m^6^A writers working via YTHDF, but not YTHDC, comprising the first phenotypes assigned to *Drosophila* mutants of the cytoplasmic m^6^A reader. These factors promote memory via neural-autonomous activities, and are required in the mushroom body, the center for associative learning. To inform their basis, we mapped m^6^A from wild-type and *mettl3* null mutant heads, allowing robust discrimination of Mettl3-dependent m^6^A sites. In contrast to mammalian m^6^A, which is predominant in 3’ UTRs, *Drosophila* m^6^A is highly enriched in 5’ UTRs and occurs in an adenosine-rich context. Genomic analyses demonstrate that *Drosophila* m^6^A does not directionally affect RNA stability, but is preferentially deposited on genes with low translational efficiency. However, functional tests indicate a role for m^6^A in translational activation, since we observe reduced nascent protein synthesis in *mettl3-KO* cells. Finally, we show that ectopic YTHDF can increase m^6^A target reporter output in an m^6^A-binding dependent manner, and that this activity is required for *in vivo* neural function of YTHDF in memory. Altogether, we provide the first tissue-specific m^6^A maps in this model organism and reveal selective behavioral and translational defects for m^6^A/YTHDF mutants.

## Introduction

Many classes of structural non-coding RNAs are heavily modified, and >150 distinct RNA modifications have been documented, e.g. on tRNA, snoRNA, snRNA, and rRNA (Boccaletto et al., 2018). These modified bases are important for normal function of these non-coding RNAs, to promote their stable secondary or tertiary structures, accumulation, and/or base-pairing interactions with trans-encoded RNA species. Many of these are constitutive modifications, although a few are now known to be reversible, raising the possibility of dynamic regulation. On messenger RNAs (mRNAs), only a limited number of modifications are known to occur; these constitute an additional layer of genetic information sometimes referred to as the epitranscriptome. Some modifications occur at the cap, and others occur at internal positions of mRNA. Although these collectively comprise a minority of nucleobases, the most prominent internal mRNA modifications include *N*^6^-methyladenosine (m^6^A), *N*^6^,2’-*O*-dimethyladenosine (m^6^Am), *N*^1^-methyladenosine (m^1^A), 5-methylcytosine (m^5^C), pseudouridine (ψ), and 2’-O-methylation (2’O-Me) (Roundtree et al., 2017a; Zhao et al., 2017a). Of these, the most abundant and most well-studied internal modification of mRNA is m^6^A (Zaccara et al., 2019).

m^6^A has been recognized to exist in mRNA since the 1970s (Desrosiers et al., 1974; Perry and Kelly, 1974), but its functional significance has been elusive until recently. Key advances that enabled its elucidation include: (1) techniques to determine individual methylated transcripts, and in particular specific methylated sites, and (2) mechanistic knowledge of factors that install m^6^A (“writers”) and mediate their regulatory consequences (“readers”). The core m^6^A methytransferase complex acting on mRNA consists of the Mettl3 catalytic subunit and its heterodimeric partner Mettl14. These associate with other proteins that play broader roles in splicing, mRNA processing and gene regulation, but that are collectively required for normal accumulation of m^6^A. It is believed that some of the roles of this larger writer complex, which includes WTAP [Fl(2)D], VIRMA (Virilizer), RBM15 (Spenito), ZC3H13, and Hakai/CBLL1, may be to recruit Mettl3/Mettl14 to chromatin and/or to specific sites in the transcriptome for modification (Zaccara et al., 2019). In addition, there is evidence for reversibility of m^6^A modifications since “eraser” enzymes such as FTO and ALKBH5 have biochemical activities to remove this mark (Roundtree et al., 2017a), although FTO has been indicated as an m^6^Am demethylase (Mauer et al., 2017).

Downstream of the writers, various readers are sensitive to the presence or absence of m^6^A, and thereby mediate differential regulation by this mRNA modification (Patil et al., 2018). The most well-characterized readers contain YTH domains, for which atomic insights reveal how a tryptophan-lined pocket selectively binds methylated adenosine and discriminates against unmodified adenosine (Luo and Tong, 2014; Theler et al., 2014; Xu et al., 2015; Xu et al., 2014). In addition, other proteins have been proposed as m^6^A readers, based primarily on preferential *in vitro* binding to methylated vs. unmethylated RNA probes. However, as adenosine methylation can affect RNA structure, care must be taken in interpreting such differential association experiments.

The functional readouts of m^6^A via different reader proteins have proven diverse and, at times, seemingly contradictory. Early studies suggested that m^6^A-containing transcripts might be unstable (Sommer et al., 1978), although without perturbations this remained correlative. However, *mettl3* loss-of-function indeed causes a global increase in m^6^A target stability (Herzog et al., 2017; Ke et al., 2017; Wang et al., 2014b). Focusing on YTH domain proteins, these fall into nuclear (DC) and cytoplasmic (DF) subclasses. The broadly expressed nuclear reader YTHDC1 has been linked to splicing (Xiao et al., 2016) and nuclear export (Roundtree et al., 2017b), while the testis-enriched nuclear reader YTHDC2 has a unique domain structure and regulates meiotic genes (Patil et al., 2018). The reader YTHDF2 destabilizes m^6^A targets (Wang et al., 2014a), by recruiting the CCR4-NOT deadenylase complex (Du et al., 2016). There are three mammalian YTHDF family members, and some evidence points to similar localization and activities of YTHDF1-3 in RNA decay and/or partitioning methylated transcripts to stress granules for silencing (Du et al., 2016; Kennedy et al., 2016; Ries et al., 2019). On the other hand, there are also reports for activating roles of DF readers. For example, YTHDF1 was cited as a translational activator (Shi et al., 2018; Wang et al., 2015), and YTHDF3 as both an RNA decay and translational stimulation factor (Shi et al., 2017). The reasons for the discrepancies remain to be resolved.

Much of our understanding of the m^6^A pathway has focused on mechanisms and genomics, but biological insights using *in vivo* genetic models have only started to emerge. Given that m^6^A mapping efforts documented thousands of modified transcripts and 10,000s of individual methylated sites across development, cell types, tissues, environmental perturbations, and diverse disease states (Liu et al., 2020; Roundtree et al., 2017a; Xiao et al., 2019; Zaccara et al., 2019), one might imagine profound phenotypic consequences to knockouts of core m^6^A pathway members. There are indeed phenotypes, but in many respects they are subtler compared to many developmental pathway mutants or other core gene regulatory factors. However, some themes have emerged in the past few years from genetic analyses.

First, there is an abundance of phenotypes in settings involving cell state transitions, such as the maternal-to-zygotic transition (Ivanova et al., 2017; Zhao et al., 2017b), or between stem cell renewal and differentiation states (Bertero et al., 2018; Vu et al., 2017; Zhang et al., 2017). This may reflect intense interest particularly in stem cell research, but on the other hand, may reflect frequent roles for RNA methylation in facilitating turnover of transcriptome programs between state or identity changes (Roundtree et al., 2017a). Second, many studies have revealed sensitivity of the mammalian nervous system to manipulation of m^6^A factors (Du et al., 2019; Widagdo and Anggono, 2018). Mutants in writer (*mettl3* and *mettl14*), reader (primarily *ythdf1*), and eraser (*FTO*) factors have collectively been shown to exhibit aberrant neurogenesis and/or differentiation (Wang et al., 2018a; Yoon et al., 2017) (Li et al., 2018; Ma et al., 2018; Wang et al., 2018b; Weng et al., 2018), which impact neural function and organismal behavior (Hess et al., 2013; Koranda et al., 2018; Shi et al., 2018; Widagdo et al., 2016). Overall, these observations may be taken as indications for the general usage of m^6^A to guide cell fate transitions along cell lineages, but may also reflect some heightened requirements in neurons, perhaps owing to their unique architectures or regulatory needs (Du et al., 2019; Livneh et al., 2019; Widagdo and Anggono, 2018).

Most insights of metazoan m^6^A biology have come from vertebrate species. Amongst invertebrates, nematodes appear to lack the core m^6^A machinery (Dezi et al., 2016), but the presence of a *Drosophila* ortholog of Mettl3 (originally referred to as IME4) opened this model system (Hongay and Orr-Weaver, 2011). While mammals contain multiple members of both nuclear and cytoplasmic YTH domain families, the fly system is simplified in containing only one of each, referred to as YTHDC (YT-521B or CG12076) and YTHDF (CG6422), respectively.

Recently, the Soller, Roignant and Lai labs established biochemical, genetic, and genomic foundations for studying the m^6^A pathway in *Drosophila* (Haussmann et al., 2016; Kan et al., 2017; Lence et al., 2016). Surprisingly, these studies jointly reported that knockout of all core m^6^A writer factors in *Drosophila* is compatible with viability and largely normal exterior patterning. Nevertheless, mutants of *mettl3*, *mettl14*, and *ythdc* exhibit a common suite of molecular and phenotypic defects. These include behavioral abnormalities as well as aberrant splicing of the master female sex determination factor *Sex lethal* (*Sxl*). Additional lines of evidence established that a critical *in vivo* function of the *Drosophila* m^6^A is to control *Sxl* splicing: (1) dose-sensitive interactions of m^6^A writer/*ythdc* mutants with *Sxl* pathway mutants, yielding enhanced female lethality, (2) detection of m^6^A at the alternatively sex-specifically spliced exon of *Sxl*, (3) sufficiency of ectopic YTHDC in male cells to induce female-specific Sxl splicing. Thus, *Sxl* splicing control is a critical *in vivo* function of the *Drosophila* m^6^A pathway.

However, major open questions from these studies, concern the regulatory and biological roles of the sole *Drosophila* cytoplasmic YTH factor, YTHDF. In contrast to other core m^6^A factors, we did not previously observe overt defects in our *ythdf* mutants, nor did it seem to exhibit robust m^6^A-specific binding activity (Kan et al., 2017). Proteomic analyses reveal YTHDC and YTHDF as the major m^6^A-specific binders in *Drosophila*, and focused biochemical tests show that YTHDF prefers a distinct sequence context than tested previously. We hypothesized that the nervous system might exhibit particular needs for the m^6^A pathway, and utilized a paradigm of aversive olfactory conditioning to reveal an m^6^A/YTHDF pathway that is important for short term memory in older animals. We complement these phenotypic data with high stringency maps of methylated transcript sites from fly heads, and show that m^6^A does not impact transcript levels but is preferentially deposited on genes with lower translational efficiency. Nevertheless, functional tests reveal that Mettl3/YTHDF can enhance translation. Finally, we show that m^6^A-binding capacity of YTHDF in neurons is necessary and sufficient to mediate normal learning and memory during aging. Overall, our study provides new insights into the *in vivo* function of this mRNA modification pathway for normal behavior.

## Results

### *Drosophila* YTHDC and YTHDF bind m^6^A in A-rich contexts

In mammals, two general classes of m^6^A-binding proteins (“readers”) are recognized, based on whether they contain or lack a YTH domain (Patil et al., 2018). Although evidence has been shown for preferential association to m^6^A vs. A for non-YTH proteins, the YTH domain is the only module for which the structural basis of selective m^6^A binding is known.

The *Drosophila* genome encodes single orthologs of nuclear (YTHDC) and cytoplasmic (YTHDF) YTH factors. We previously tested capacities of their isolated YTH domains to associate preferentially with m^6^A, using RNA probes bearing GGm^6^ACU vs. GGACU contexts (Kan et al., 2017). This motif represents the favored binding site for mammalian YTHDC1, which has explicitly been shown to prefer G and disfavor A at the -1 position (Xu et al., 2014). Of note, however, mammalian YTHDF1 does not share this discriminatory feature (Xu et al., 2015). We previously observed the YTH domain of *Drosophila* YTHDC exhibits robust and selective binding to this methylated probe, but the corresponding domain of YTHDF had only modest activity. From these tests, it was not clear whether the isolated YTH domain might not be fully functional, or perhaps prefers a distinct target site. We tested both of these notions.

We compared the binding of full-length YTHDC and YTHDF proteins to m^6^A vs. A using biotinylated RNA photoaffinity probes (Arguello et al., 2017). These probes contain diazirine-modified uridine (5-DzU) that can be cross-linked to protein upon UV irradiation (**Figure 1A**). We have shown that 5-DzU does not interfere with protein binding at the modified nucleotide, and therefore enables high-efficiency detection of associated proteins (Arguello et al., 2017). We incubated cell lysates expressing tagged YTH proteins with beads conjugated to GGm^6^ACU/GGACU RNA probes, immunoprecipitated complexes with streptavidin, and performed Western blotting for YTH factors. We observed modestly enhanced association of YTHDC to GGm^6^ACU vs. GGACU, while YTHDF did not crosslink preferentially to this methylated probe (**Supplementary Figure 1**).

**Figure 1.**
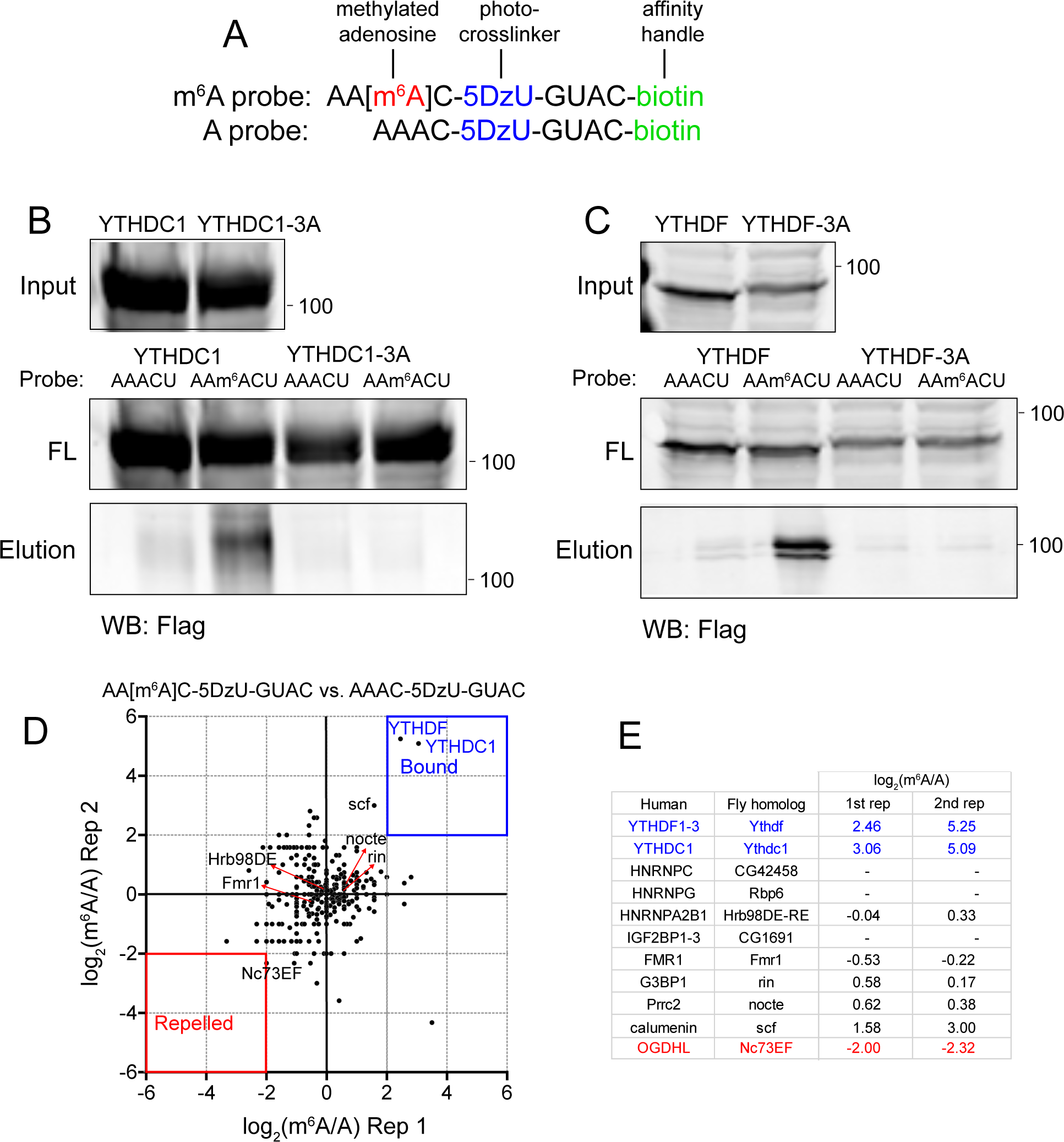
YTHDC and YTHDF are the major m^6^A-binding proteins in *Drosophila* (A) RNA photoaffinity probes used in crosslinking assays. (B-C) Both nuclear (YTHDC) and cytoplasmic (YTHDF) *Drosophila* YTH factors specifically recognize m^6^A within the AAm^6^ACU context. Point mutations of aromatic residues that line the m^6^A cage (3A variants) abolish selective binding to m^6^A probes. FT: Flow through (C) Proteomic profiling of S2 cells using m^6^A and A RNA photoaffiinity probes reveals YTHDC and YTHDF as the only preferentially bound (reader) proteins, and no strongly repelled proteins were found in these conditions. Background proteins are clustered together around the plot origin; threshold=2X the interquartile range. (D) Selected values of fly homologs of mammalian m^6^A readers and repelled proteins expressed in S2 cells, as well as candidates of novel bound/repelled factors.

As our previous mapping suggested that *Drosophila* m^6^A modifications are biased to have upstream adenosines (Kan et al., 2017), we next compared AAm^6^ACU/AAACU probes. Interestingly, both YTHDC and YTHDF exhibited clearly preferential binding to methylated adenosine in this context (**Figure 1B-C**). Next, we tested variants in which three critical tryptophan/leucine residues in the m^6^A binding pocket were mutated to alanine (**Supplementary Figure 1**). Although “3A” mutant proteins accumulated to similar levels as their wild type counterparts, both YTHDC-3A and YTHDF-3A failed to bind m^6^A (**Figure 1B-C**), indicating that their specificity for methylated RNA requires intact YTH domains. Thus, *Drosophila* YTH proteins, in particular YTHDF, may prefer an A-rich context.

### YTHDC and YTHDF are the dominant *Drosophila* m^6^A-binding proteins

Having clarified that both fly YTH factors specifically discriminate between m^6^A and A, we sought to identify differential binders using an unbiased approach. Proteomic studies in mammalian cells reveal YTH factors as dominant proteins that preferentially associate with m^6^A compared to unmethylated probes, along with some other proteins (e.g. FMR1 and LRPPRC), and reciprocally some factors that are repelled by this modification (e.g. stress granule factors such as G3BP1/2, USP10, CAPRIN1, and RBM42) (Arguello et al., 2017; Edupuganti et al., 2017). As well, other methods were used to identify mammalian factors that appear to bind preferentially to m^6^A, such as Prrc2a (Wu et al., 2019) and IGF2BP1-3 (Huang et al., 2018).

We used our AAm^6^ACU/AAACU RNA photoaffinity probes to pull down endogenous proteins from S2 cell lysates, followed by mass spectrometry. We performed replicate proteomic assays, and plotted the ratios of peptide counts recovered from m^6^A and A probes (**Figure 1D** and **Supplementary Table 1**). These experiments revealed YTHDC and YTHDF were strongly and reproducibly enriched with the m^6^A probe compared to the A probe. By contrast, we did not observe clearly differential association of any other factors, including all fly homologs of other mammalian proteins reported to preferentially bind or be repelled by m^6^A (Patil et al., 2018) (**Figure 1D-E**).

Because several recent studies examined the relationship of mammalian FMRP as an m^6^A binding protein (Arguello et al., 2017; Edupuganti et al., 2017), and subsequently as an m^6^A effector (Edens et al., 2019; Hsu et al., 2019; Zhang et al., 2018a), we tested further if we could relate *Drosophila* FMR1 to m^6^A. In our tests, FMR1 was actually mildly depleted from m^6^A probe relative to the A probe (**Figure 1C-D**), although peptide counts were similar in both cases (**Supplementary Table 1**). We tested further if FMR1 might associate with the demonstrated YTH readers. Using tagged constructs, we found that FMR1 and YTHDF could be reciprocally co-immunoprecipitated in S2 cells (**Supplementary Figure 2**). A potential hint for direct association came with the observation that YTHDF-3A also interacted with FMR1 in reciprocal co-IP tests. However, all of these interactions were largely eliminated upon treatment of lysates with RNase I (**Supplementary Figure 2**), suggesting that they were bridged by RNA.

Overall, while it is possible that other target sequences or lysate sources might reveal other differential binders, these analyses led us to focus on YTH domain factors as the major direct readers for m^6^A biology in *Drosophila*.

### Neural autonomous function of m^6^A supports olfactory learning

The expression of several m^6^A factors is elevated in the *Drosophila* nervous system, and mutants of several m^6^A factors are viable, but exhibit similar locomotor defects (Haussmann et al., 2016; Kan et al., 2017; Lence et al., 2016). As this suggested preferential sensitivity of the nervous system to m^6^A, we examined phenotypic requirements of neural m^6^A in greater detail.

Recent studies showed that m^6^A pathway is required for learning and memory in mice (Shi et al., 2018; Zhang et al., 2018c). We used a classical aversive conditioning paradigm to test *Drosophila* m^6^A mutants for deficits in short-term memory (henceforth ‘STM’ or ‘memory’). To obtain time resolved performance measurements, we employed a conditioning apparatus (multifly olfactory trainer–MOT, **Figure 2A** and **Supplementary Figure 3**) (Claridge-Chang et al., 2009; Tumkaya et al., 2018). Briefly, one odor is administered in the presence of a shock stimulus while the other odor is subsequently delivered in the absence of foot shock (**Figure 2B**). Because shock is innately aversive, *Drosophila* will associate the odor given in the presence of shock with harm and will tend to avoid it during subsequent encounters. During the test phase, flies are presented with both odors; the avoidance of the conditioned odor can be quantified to measure an aversive olfactory memory, which can last up to two hours for a single conditioning assay.

**Figure 2.**
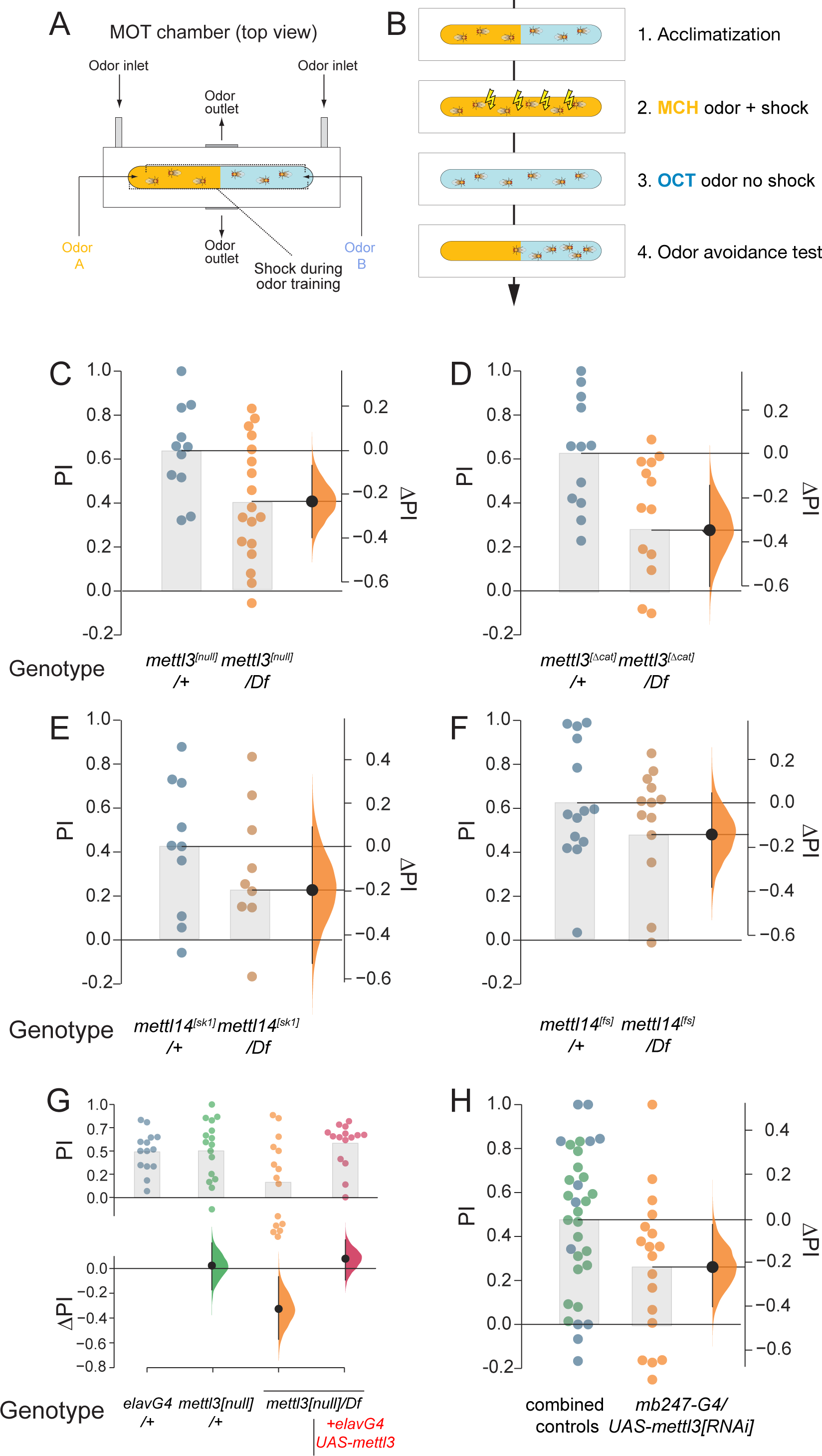
m^6^A pathway is required for short term learning and memory (STM) in *Drosophila* (A) Schematic of the multifly olfactory training (MOT) chamber apparatus used for behavioral measurements. (B) Paradigm for shock-associated odor avoidance assay for STM acquisition. All assays in C-H were conducted in 20 day old flies. (C-D) Hemizygote null conditions for *mettl3[null]* and *mettl3[Δcat]* both led to (≈Δ PI-0.2) STM impairment. (C-D) Hemizygote null conditions for *mettl14[fs]* and *mettl14[sk1]* led to a moderate (≈Δ PI-0.2) STM impairment. (E) Pan-neuronal expression of *UAS-mettl3* using *elav-Gal4 (elav-G4)* in *mettl3[null]* hemizygotes rescued their STM phenotype. (F) Celltype-specific knockdown of *mettl3* in the mushroom body using *mb247-Gal4* phenocopied STM impairment seen in whole animal *mettl3* mutants.

We did not observe any memory impairment in heterozygous m^6^A LOF mutants (data not shown), and therefore used the respective heterozygotes as controls in subsequent tests. To minimize background genetic effects, which are a frequent confound of behavioral assays, we compared these to trans-heterozygous or hemizygous (over deficiency) allelic combinations. In young flies, both writer mutants were essentially normal: we observed only a modest memory reduction in 10-day-old *mettl3* hemizygous nulls, while similarly aged *mettl14* mutants showed no impairment (**Supplementary Figure 4)**. However, at 20 days, both *mettl3[null]* and *mettl3[Δcat]* hemizygous nulls displayed a substantially stronger (ΔPI -0.2 to -0.3) STM impairment (**Figure 2C-D**). Consistent with the role of Mettl14 as a cofactor for Mettl3, hemizygote *mettl14[fs]* and *mettl14[SK1]* mutants also exhibited comparable memory impairments in 20-day-old flies (**Figure 2E-F**).

Assays of whole animal mutants did not resolve if the nervous system *per se* was involved in these behavioral defects. We addressed this using tissue-specific knockdown and rescue experiments. We first generated *mettl3[null]* hemizygote animals bearing *elav-Gal4* and *UAS*-*mettl3* transgenes, to drive their expression in all neurons. In this genetic background, all non-neuronal cells of the intact animal lack Mettl3. Strikingly, these flies exhibited normal memory (**Figure 2G**), providing stringent evidence that the odor avoidance behavioral defect of m^6^A knockouts is strictly due to a cell-autonomous function of Mettl3 in neurons.

In *Drosophila, a*ssociative olfactory memory is formed within the mushroom body (de Belle and Heisenberg, 1994; Heisenberg et al., 1985). To test whether m^6^A is specifically required in the mushroom body, we validated the ability of a *UAS-mettl3[RNAi]* transgene to deplete Mettl3 protein (**Supplementary Figure 5A**), and applied *mettl3* knockdown to a majority of mushroom-body neurons using *MB247-Gal4*. This manipulation impaired memory to a degree that was comparable to the impairment in whole animal *mettl3* null mutants (**Figure 2H**). Therefore, m^6^A is specifically required in the *Drosophila* mushroom body to mediate odor avoidance learning.

### YTHDF, but not YTHDC, is the functional effector of m^6^A during STM

We sought to elaborate the regulatory pathway underlying m^6^A in learning and memory. Prior genetic assays linked m^6^A writers Mettl3/Mettl14 in a pathway with nuclear reader YTHDC for locomotor and gravitaxis behaviors, as well as ovary development (Haussmann et al., 2016; Kan et al., 2017; Lence et al., 2016). By contrast, our *ythdf* mutants did not resemble other core m^6^A mutants, and overall seemed to lack substantial defects in these assays (Kan et al., 2017).

The phenotypic discrepancy of these mutants was further emphasized by quantifying their lifespans. While mutations in *mettl3* and *ythdc* led to severely shortened lifespan (>40 days), loss of *ythdf* decreased lifespan only modestly (**Figure 3A-C** and **Supplementary Figure 6A-C**). As some behavioral effects of the m^6^A pathway are mediated by the nervous system (Lence et al., 2016), we tested the effect of pan-neuronal depletion of *mettl3* using *elav-Gal4*.

**Figure 3.**
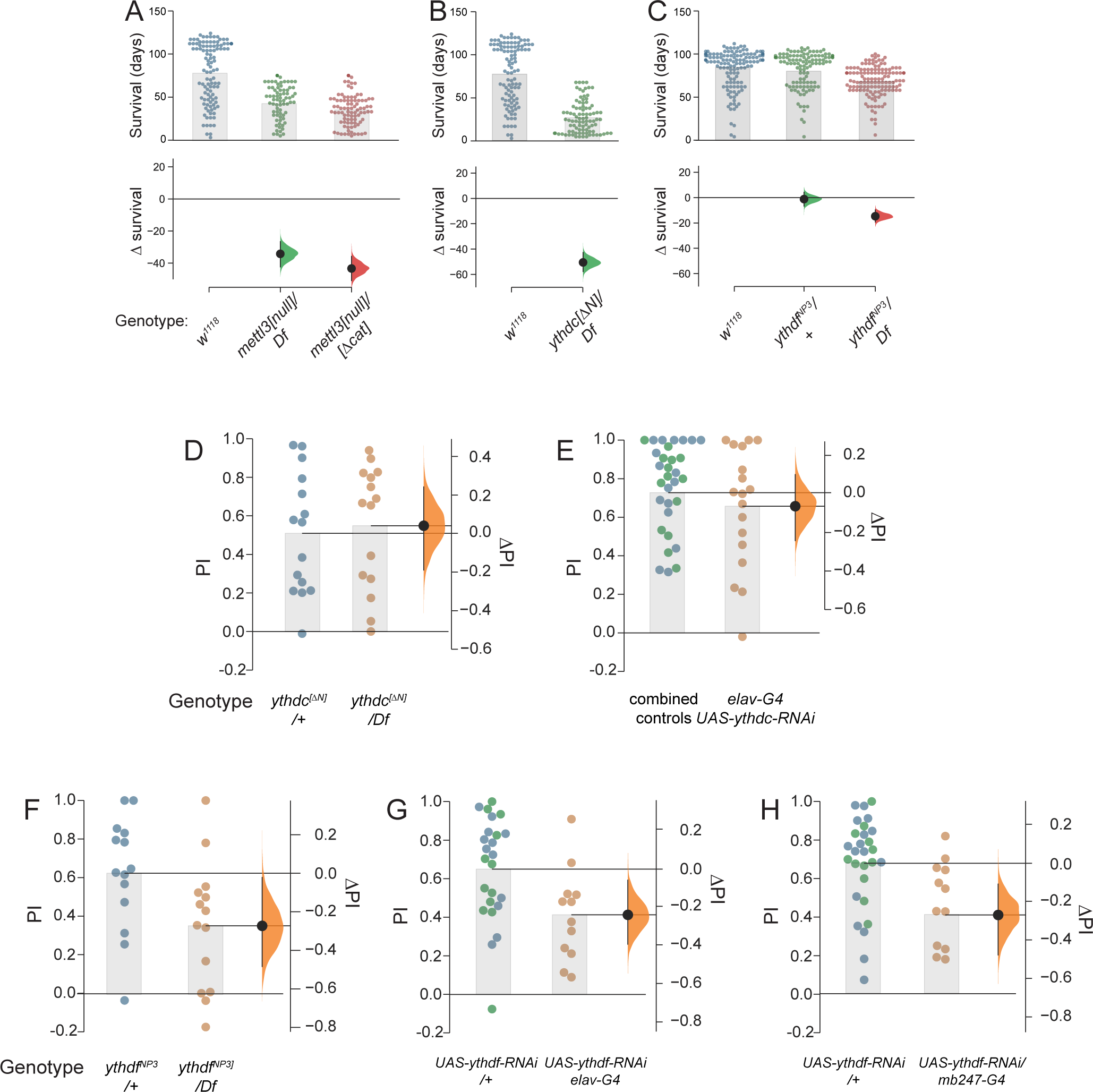
YTHDF, but not YTHDC, mediates the role of m^6^A in *Drosophila* STM (A-C) Lifespan measurements of m^6^A writer and reader mutants. Mutants of *mettl3* (A) and nuclear reader *ythdc* (B) exhibit severely shortened lifespan, but mutants of cytoplasmic reader *ythdf* (C) have only minor defect in lifespan. (D-I) STM measurements in 20 day flies. (D) Despite gross behavioral defects and short lifespan, *ythdc* mutants exhibit normal STM. (E) Pan-neuronal knockdown of *ythdc* using *elav-Gal4* (*elav-G4*) also yields normal STM. (F) *ythdf* hemizygotes recapitulate age-induced STM impairment seen in m^6^A writer mutants. (G-H) Pan-neuronal knockdown (G) or mushroom body-specific knockdown (H) of *ythdf* compromised STM similar to whole animal *ythdf* mutants.

Compared to controls, lifespan was modestly (10 days) shorter (**Supplementary Figure 6D**). These data support the concept of a physiologically important role for nuclear readout of m^6^A *via* the YTHDC reader, with major, specific nervous-system effects on longevity.

In light of extensive locomotor defects and short lifespan of *ythdc* mutants, we were surprised to find that they lacked memory deficits at either 10 or 20 days of age (**Supplementary Figure 4** and **Figure 3D**). We also tested cell-specific depletion of *ythdc* using RNAi (**Supplementary Figure 5B**); pan-neuronal knockdown of *ythdc* using *elav-Gal4* in aged flies also did not affect STM (**Figure 3E**). Thus, we were prompted to examine mutants of the cytoplasmic reader YTHDF more carefully. Excitingly, *ythdf* hemizygotes exhibited age-related memory impairment (**Figure 3F**), comparable to *mettl3* mutants. Since *ythdc* mutants generally phenocopy other defects of m^6^A writer mutants, these data indicate a division of labor between the *Drosophila* YTH readers, downstream of m^6^A writers.

To test whether YTHDF was specifically required in the nervous system and/or the MB, we used a validated RNAi transgene to deplete it in a tissue-specific manner (**Supplementary Figure 5B**). Upon knockdown of *ythdf* with either *elav-Gal4* or *mb247-Gal4*, 20-day-old flies exhibit impaired memory (**Figure 3G-H**). Altogether, these data indicate that cytoplasmic readout of m^6^A by YTHDF is required in older flies for the normal functioning of memory-storing neurons.

### Neither Mettl3 nor YTHDF can cross-rescue each other’s memory defects

We next asked whether overexpression of YTHDF in *mettl3* mutants, or the reciprocal genetic manipulation, would affect memory. Successful rescue could, for example, suggest that the reading function of YTHDF is not fully dependent on Mettl3 methylation, i.e. may somehow involve a parallel pathway. However, in *mettl3* nulls supplemented with pan-neuronal YTHDF overexpression had no memory improvement (**Supplementary Figure 7A**). Similarly, overexpression of Mettl3 did not alter improve the memory impairment of *ythdf* mutants (**Supplementary Figure 7B**). Beyond serving as stringent negative controls then, for the cognate rescue experiments (**Figures 2 and 3**), these results provide further credence to the notion that a linear, directional Mettl3/4 → YTHDF pathway underlies m^6^A-mediated memory function.

### Mapping the Mettl3-dependent m^6^A methylome in *Drosophila*

To link these brain-function defects to the underlying molecular landscape of RNA methylation, we sequenced m^6^A sites from polyadenylated transcripts using miCLIP (Linder et al., 2015). Although we previously reported miCLIP datasets from *Drosophila* embryos (Kan et al., 2017), we recognized that there can be background association in such data. Thus, individual sequencing “peaks” need to be interpreted cautiously. To provide a stringent basis to infer the existence of m^6^A at given sites, we analyzed companion input and miCLIP libraries from dissected heads, which are highly enriched for neurons, comparing wild-type and deletion mutants of *mettl3*, which encodes the catalytic methyltransferase subunit essential for mRNA modification (e.g. **Figure 1** and **Supplementary Table 2**).

The miCLIP libraries from *mettl3* mutants proved especially valuable, because they allowed us to distinguish m^6^A-IP loci that were clearly genetically dependent on endogenous Mettl3 (**Figure 4A**, **Supplementary Figure 8A**). Reciprocally, numerous regions of the transcriptome were significantly enriched in miCLIP libraries compared to input, but whose signals persisted in *mettl3* mutants (**Figure 4B, Supplementary Figure 8B**). These might conceivably represent transcript regions modified by another factor (Pendleton et al., 2017), but cannot at this point be easily distinguished from non-specific pulldown. In general, the Mettl3-independent peaks were globally present in weaker m^6^A peaks (**Figure 4C**), suggesting they are functionally less relevant. Therefore, we applied stringent filtering to focus our attention on the rich set of clearly Mettl3-dependent peaks (**Figure 4A-C**). In addition, as we employed strong selection for polyadenylated transcripts for input, we prioritized studies of annotated genes. Altogether, our analyses (see Methods) yielded 3874 Mettl3-dependent peaks from 1635 genes. Since a subset of these called regions contained clear local minima, we applied PeakSplitter (Salmon-Divon et al., 2010) to arrive at 4686 head m^6^A peaks (**Supplementary Table 3**).

**Figure 4.**
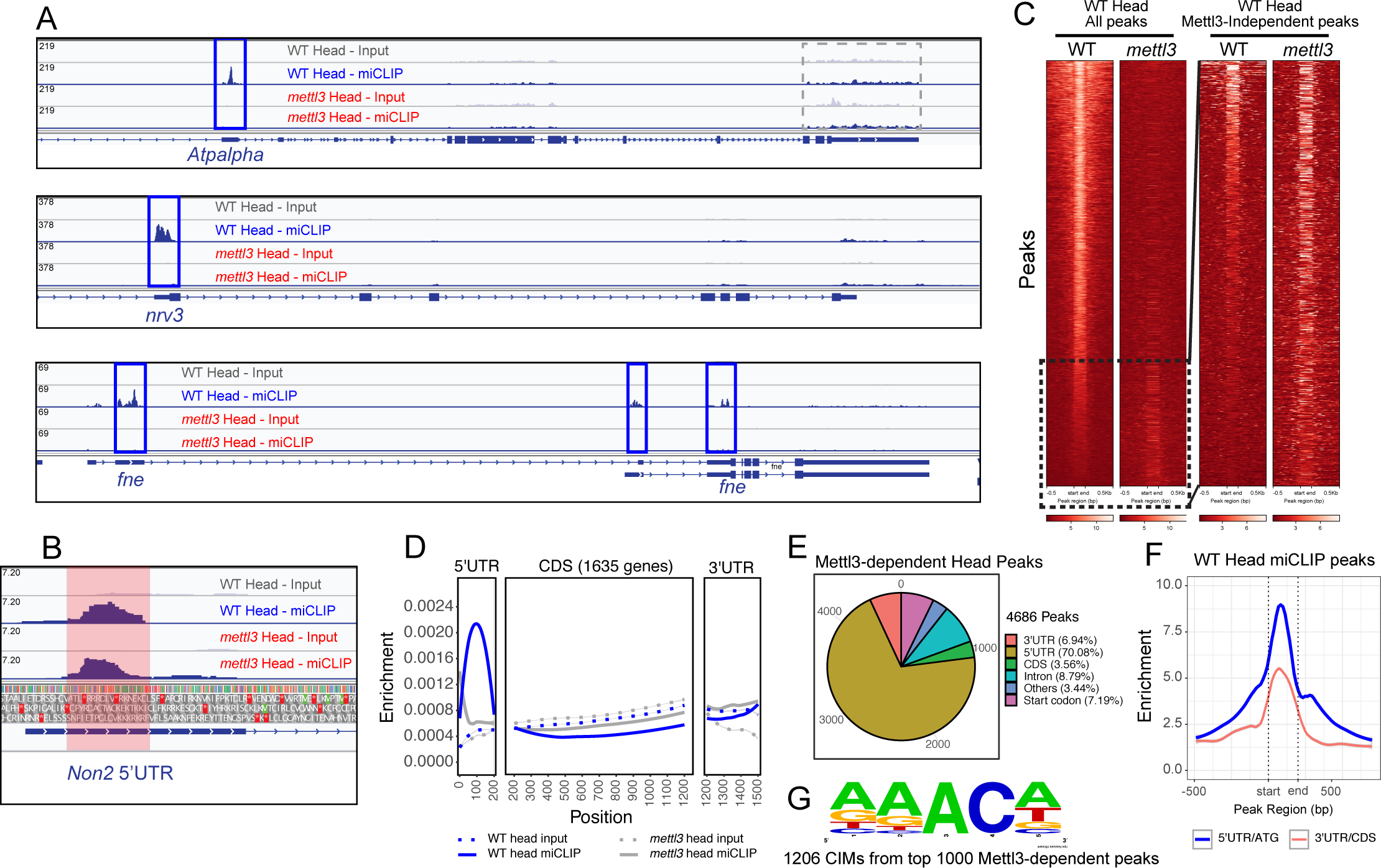
High stringency mapping of the *Drosophila* m^6^A methylome reveals new features (A) IGV screenshots of genes that exemplify archetypal 5’ UTR miCLIP enrichment and the utility of wild type vs. *mettl3* mutant comparisons. The IGV tracks above the transcript model depict miCLIP and input libraries from wild type and *mettl3* female heads. Note the 5’ UTRs of all three genes (*Atpalpha, nrv3* and *fne)* contain prominent Mettl3-dependent peaks (blue box). Other exonic regions sequenced in miCLIP libraries are not enriched above input libaries (e.g. grey box). (B) IGV screenshot of *Non2* illustrates a 5’UTR Mettl3-independent peak (red box). (C) Heatmaps of Mettl3-dependent and -independent m^6^A peaks. Heatmaps of input normalized miCLIP signals at Mettl3-dependent and independent peaks. Each line represents miCLIP enrichment over input across a MACS2-called peak, as well as 500nt flanking regions. Heatmaps on the left are all peaks including Mettl3-dependent and independent m^6^A peaks. Mettl3-independent m^6^A peaks are displayed in the right panels. Note that the scales are different for the heatmap panels. Approximate locations of Mettl3-independent m^6^A peaks within all peaks are indicated by a dashed box. (D) Metagene profiles of miCLIP and input signals along a normalized transcript using genes that contain high-confidence m^6^A peaks in head libraries. An overwhelming 5’ UTR enrichment is observed. (E) Metagenes of enrichment at high-confidence m6A peaks that have been group as 5’ UTR/start codon and 3’ UTR/CDS regions. Metagenes are produced by averaging signals from input normalized miCLIP from WT head. On average, stronger signals are observed at 5’ UTRs/start codons. Peak start and end are specified on the x-axis. Dashed lines include the start and end locations of peaks. (F) Pie chart depicting the fraction of m^6^A peaks in different transcript segments. (G) Nucleotide content surrounding CIMs located within the top 1000 Mettl3-dependent m^6^A peaks in head.

### *Drosophila* m^6^A is highly enriched in 5’ UTRs within adenosine-rich contexts

Characterization of *Drosophila* Mettl3-dependent m^6^A peaks revealed fundamental similarities and differences with m^6^A patterns in other organisms. Mammalian (e.g. human and mouse) m^6^A is well-known to dominate at stop codons and 3’ UTRs (Zaccara et al., 2019; Zhao et al., 2017a). In fish, m^6^A is also highly enriched at stop codons, but the predominant Mettl3-dependent signals localize to 5’ UTRs (Zhang et al., 2017). Previous work in *Drosophila* was conflicting, since low-resolution meRIP-seq suggested mostly CDS modification with a small minority in UTRs (Lence et al., 2016), while our prior miCLIP data indicate dominant UTR modifications, preferentially in 5’ UTRs (Kan et al., 2017). However, these maps were generated with different technologies, and neither was controlled against mutants.

Our new miCLIP data provide a clearer perspective. Strikingly, while found at some level throughout the transcriptome, m^6^A predominates in 5’ UTRs in *Drosophila*. This can be observed at numerous individual loci (**Figure 4A** and **Supplementary Figure 8**) and via miCLIP metagene profiles (**Figure 4D**). Overall, while we do observe some Mettl3-dependent coding sequence (CDS) and 3’ UTR miCLIP peaks (**Figure 4E, and Supplementary Figure 8**), these were overall rare, of generally lower ranks than 5’ UTR and start codon peaks (**Figure 4F**), and not appreciably enriched in metagene profiles over companion mutant datasets (**Figure 4D**).

We examined C-to-T crosslinking-induced mutations following adenosine residues (CIMs), which have been taken to represent individual m^6^A site in miCLIP data (Grozhik et al., 2017; Linder et al., 2015). In particular, we focused on CIMs located within Mettl3-dependent m^6^A-IP peaks, which we took as bearing high-confidence RNA methylation sites. Within these, the sequence context of CIMs in *Drosophila* roughly resembles the DRAC context that has been observed in other species (Zaccara et al., 2019). However, while a majority of sites fall into a GGACH context in vertebrates (Zaccara et al., 2019), m^6^A sites in *Drosophila* prefer AAACD (**Figure 4G**), correlating with the preferred binding sites of YTHDC and YTHDF in our assays of photocrosslinking-activated m^6^A probes (**Figure 1**).

We validated our map by testing m^6^A-IP to IgG-IP samples for enrichment of m^6^A target transcripts using rt-qPCR (**Supplementary Figure 9**). We validated a number of top m^6^A targets (e.g. *aqz*, *Syx1A*, *fl(2)d*, *Prosap*, *pum, futsch*, *gish*) from whole female fly RNA (**Figure 5A**). Still, recognizing that m^6^A-RIP-qPCR evaluates the presence of entire transcripts in pulldowns, we performed parallel experiments from *mettl3[null]* female flies. All of these binding events, even loci with very modest enrichment in wild type (e.g. *sky*, **Figure 5A**), were found to be Mettl3-dependent. By contrast, control loci lacking m^6^A peaks (*fwe* and *CG7970*) showed very little m^6^A-dependent IP signals, and these were unaltered in *mettl3[null]* samples (**Figure 5A’**). These data provide stringent validation of our m^6^A maps.

**Figure 5.**
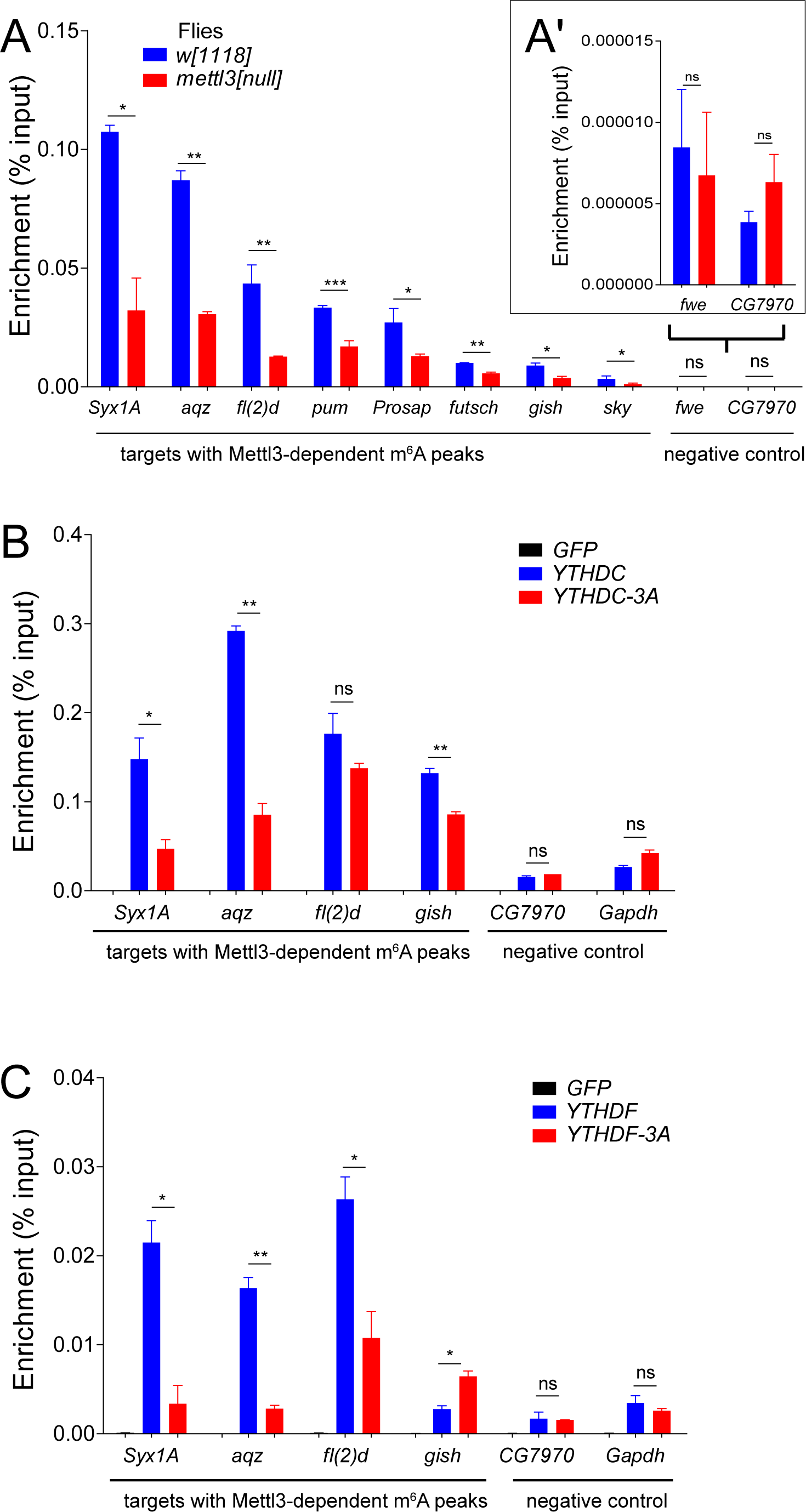
Validation of *Drosophila* m^6^A targets and their association with readers (A-A’) m^6^A-RIP-qPCR validation of m^6^A transcripts in *w[1118]* control and *mettl3[null]* knockout whole female flies. (B-C) RIP-qPCR of m^6^A targets in S2-S cells and transfected YTHDC (C) or YTHDF (C) constructs shows specific pulldown of several targets relative to GFP control, and the association of several of these is compromised by mutation of the YTH domain (3A versions).

Overall, our high quality miCLIP data from the *Drosophila* head reveals that the position of m^6^A in this species appears distinct amongst metazoans (highly 5’ UTR specific) and occurs within a distinct adenosine-rich context.

### *Drosophila* YTH factors associate with m^6^A targets in a YTH-dependent manner

We next assessed association of m^6^A targets with YTH factors using transfected constructs in S2-S cells. Although overexpression may affect the localization properties of YTH domain proteins (Ries et al., 2019), we showed that ectopic YTHDC and YTHDF localize to the nucleus and cytoplasm of cultured cells, respectively (Kan et al., 2017). In these tests, it is also relevant to consider that we are evaluating the association of the test proteins and target RNAs, which may or may not occur directly through the modified nucleotides. However, we can compare these to YTH-“3A” point-mutant counterparts that disrupt m^6^A selectivity (**Figure 1**).

We immunoprecipitated tagged YTH wild-type or “3A” mutant factors and performed qPCR for validated m^6^A targets or negative control transcripts. By comparison to control GFP-IP, we observed preferential binding of YTHDC/YTHDF on multiple m^6^A targets, compared to non-m^6^A transcripts (**Figure 5B-C**). By testing companion “3A” mutant factors, we gained evidence for direct association of YTH factors on m^6^A targets. However, a clear picture of target selectivity did not emerge (**Figure 5B-C**). *Syx1A* and *aqz* exhibited the most clearly differential association between wt and 3A forms of both YTHDC and YTHDF. We observed potentially selective association with other loci, in that *gish* was preferentially bound only by wt YTHDC while *fl(2)d* was preferentially bound only by wt YTHDF.

We bear in mind these were ectopic experiments, and thus cannot rule out non-physiological associations. Even though we observed many cases of YTH-dependent target association, both YTH-3A proteins still exhibited apparent enrichment compared to GFP. If these mutant YTH proteins are still capable of incorporating into RNA granules, this may conceivably indicate indirect interactions with transcripts. Nevertheless, these data provide evidence that YTH domain proteins, including YTHDF, associate with specific m^6^A target transcripts via their m^6^A-binding pocket in *Drosophila* cells.

### m^6^A does not globally influence mRNA levels in *Drosophila*

There is diverse literature on linking mammalian YTHDF homologs to RNA decay and/or translational activation, while the function of *Drosophila* m^6^A/YTHDF has been little studied. The only prior study integrated MeRIP-seq peaks from S2R+ cells with RNA-seq data from m^6^A pathway depletions, and concluded that m^6^A exerts a slight positive influence on mRNA levels (Lence et al., 2016). With our high-stringency m^6^A map from heads, we generated RNA-seq data from one- and three-week old heads using *mettl3*, *ythdf* heterozygotes and transheterozygotes. The heterozygote samples provide matched genetic backgrounds for comparison, and the temporal series assesses CNS stages including an advanced setting during which behavioral phenotypes were apparent (**Figures 2-3**).

Transcriptome analyses revealed scores of differentially expressed genes in one- and three-week old mutants (**Supplementary Table 5)**, a majority of which were uniquely misexpressed (**Supplementary Figure 10A-B**). Most affected genes were not found to be common between m^6^A writer (*mettl3*) and reader (*ythdf*) mutants, although there were mild changes that gradually increased with tissue age (**Supplementary Figure 10C and D**). Thus, there did not appear to be a clear signature of m^6^A/YTHDF regulation revealed by bulk gene expression.

We examined this more closely by directly examining the behavior of m^6^A targets. We reasoned that targets with systematically higher levels of methylation - that is, genes with increasing proportions of methylated transcripts - would be more sensitive to loss of the m^6^A pathway. However, while our miCLIP libraries provide Mettl3-dependent peaks and single nucleotide resolution mapping of m^6^A sites in the transcriptome, it is not possible to infer overall methylation levels. A solution to this limitation, grouping targets by number of sites/peaks, has been adopted by others (Cheng et al., 2019; Ries et al., 2019) and proposes that targets with increasing numbers of peaks/sites may have more individual transcripts with at least one m^6^A modification. Therefore, we binned genes by numbers of Mettl3-dependent m^6^A peaks.

In contrast to prior association of *Drosophila* m^6^A with mRNA stabilization (Lence et al., 2016), we did not observe many changes in our high-confidence m^6^A targets in *mettl3* (**Figures 6A**) or *ythdf* mutant CNS from any stage (**Supplementary Figure 10E-H**). Paradoxically, even though all bins of m^6^A targets clustered closely with a log_2_ fold change of 0, Kolmogorov-Smirnov (KS) tests indicated statistical significance when comparing sets of m^6^A targets and background. While statistically different, our analyses clearly demonstrate that there are no directional gene expression changes in methylated transcripts under writer or reader loss (**Figures 6A and Supplementary Figure 10**).

**Figure 6.**
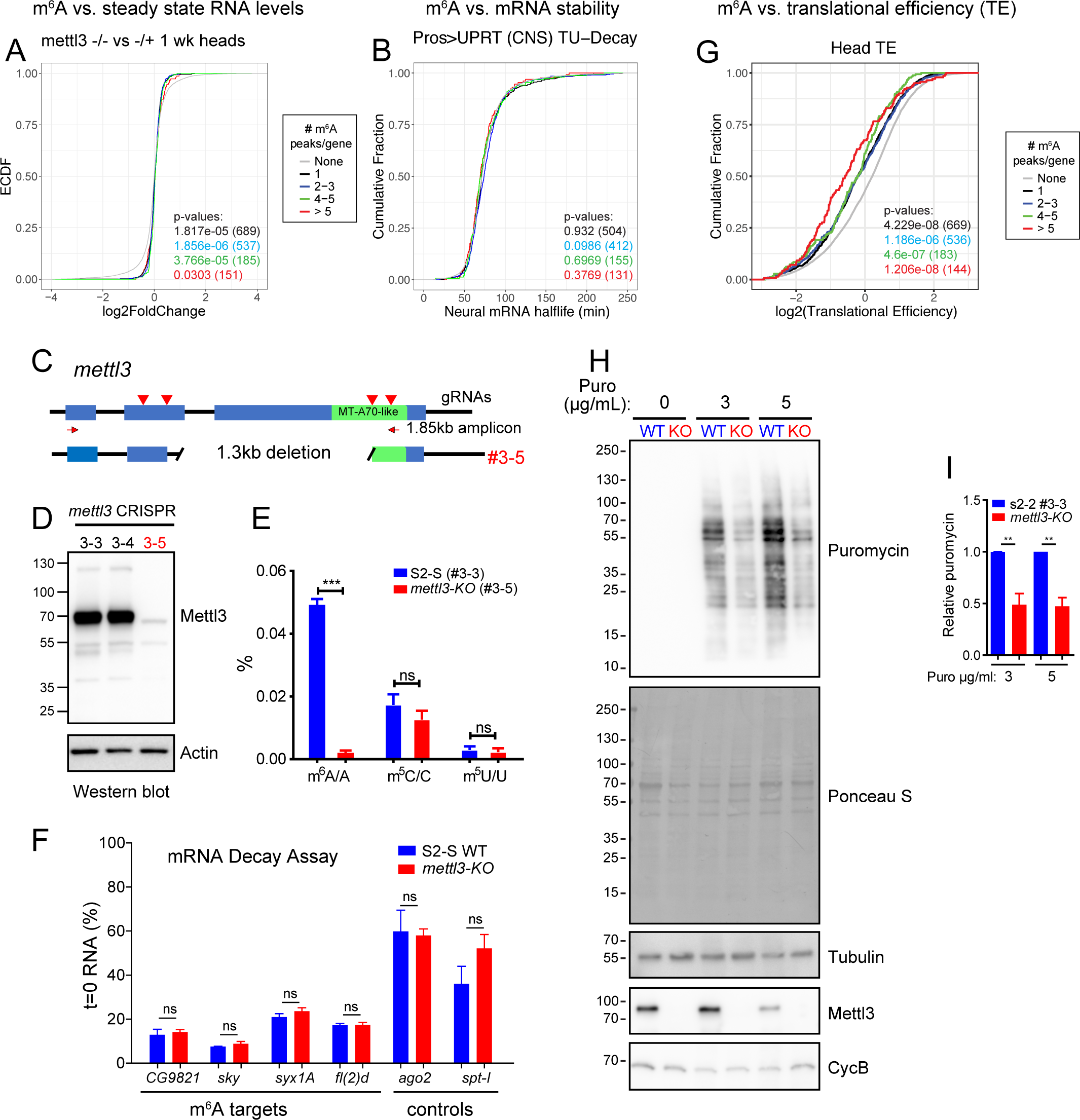
m^6^A mediates translational activation in *Drosophila* (A) Steady state RNA levels of m^6^A targets are not affected by loss of m^6^A. Differential gene expression analysis was performed comparing *mettl3* heterozygote and transheterozygote fly heads that were aged to 1 week. No directional change is observed in any group of m^6^A targets, binned by increasing numbers of Mettl3-dependent m^6^A peaks. (B) m^6^A targets are not biased in their mRNA stability. The half-life of neural genes obtained from TU-decay measurements of the nervous system (Burow et al., 2015) is plotted as cumulative distribution grouping target genes as in (A) based on number of peaks per target. (C) We used CRISPR/Cas9 to delete the *mettl3* locus from S2-S cells. Genotyping of clonal lines identifies 1.3 kb biallelic deletion in the #3-5 line. (D) Western blotting validates absence of Mettl3 protein in #3-5 (referred to as KO) line compared to other lines subjected to CRISPR; #3-3 was subsequently used as a control S2-S cell line. (E) Levels of different modified nucleosides in *mettl3* knockout cells using quantitative liquid chromatography–mass spectrometry (LC-MS) and absolute quantification confirms specific lack of m^6^A. (F) qPCR measurements of m^6^A-modified mRNAs show that five hours after actinomycin-D treatment, they have similar half-lives in *mettl3-KO* cells as in wild-type cells. NS, not significant. (G) Translation efficiency (TE) measurements (Zhang et al., 2018b) plotted as cumulative fractions for targets with different numbers of m^6^A peaks. m^6^A is preferentially deposited on genes with low TE. (H) Puromycin labeling assay shows the global protein synthesis reduced in S2-S cells lacking Mettl3 writer. (I) Quantification of reduced nascent protein synthesis in *mettl3-KO* S2-S cells. For (A-B and G) a bootstrap method generated the background distribution (None) using genes that lacked m^6^A peaks. To generate p-values, two-sided Kolmogorov-Smirnov (KS) tests were performed comparing the background distribution and each group of m6A target genes. Number of targets are included in parentheses.

Since it was conceivable that some expression trends were masked in steady state measurements, we examined a published dataset of *in vivo* mRNA decay rates generated using dynamic TU-tagging from the *Drosophila* CNS, obtained by pulse-chase labeling of *pros-Gal4>UAS-UPRT* cells with 4-thiouridine (Burow et al., 2015). We observed that, in aggregate, m^6^A-modified transcripts had identical mRNA half-lives as the background distribution (**Figure 6B**). Thus, we were not able to discern global m^6^A regulatory impacts on transcript properties.

To test this further, we used CRISPR/Cas9 to delete the *mettl3* locus from S2-S cells (**Figure 6C**), a derivative of S2 cells lacking viruses (Reimao-Pinto et al., 2016). Although we initially recovered heterogenous events, we were able to obtain clonal lines following serial dilution (**Supplementary Figure 11**). We used #3-5 bearing a 1.3 kb biallelic deletion (**Supplementary Figure 11**) and used a sibling clone #3-3 from the mutagenesis procedure that did not harbor alterations in *mettl3* as a control line. Western blotting validated absence of METTL3 protein in #3-5 (referred to as *mettl3-KO*) line compared to #3-3 (referred to as wt S2-S) cells (**Figure 6D**). Moreover, we directly measured *N*^6^-methyladenosine levels in *mettl3-KO* cells using quantitative liquid chromatography–mass spectrometry (LC-MS). External calibration curves prepared with A and m^6^A standards determined the absolute quantities of each ribonucleoside. The mRNA m^6^A methylation levels in knockout cells were decreased to <5% of that in wild-type cells, whereas other modified ribonucleosides were unaffected (**Figure 6E**).

Using the *mettl3-KO* cells, we performed RNA decay assays of validated m^6^A targets and control transcripts. Following inhibition of transcription using actinomycin D, we observed a range of transcript levels across different loci, but none of these were significantly different between wild-type and m^6^A-deficient cells (**Figure 6F**). Overall, our analyses using S2 cells and intact nervous system indicate that mRNA stability of m^6^A-containing transcripts is neither substantially nor directionally influenced by loss of m^6^A in *Drosophila*, in contrast to m^6^A in mammals.

### m^6^A is preferentially deposited on fly transcripts with lower translational efficiency

In light of these data, we examined the alternate possibility of m^6^A-dependent translational control. For this purpose, we utilized ribosome profiling datasets from *Drosophila* heads (Zhang et al., 2018b) to assess translational efficiencies of transcripts with or without m^6^A modifications. Strikingly, we found that genes with m^6^A had lower translational efficiency than the background distribution (**Figure 6G**). The functional relevance of this observation was strengthened by the fact that the number of m^6^A peaks per transcript exhibited a progressive, inverse correlation with translational efficiency and contrasted with the lack of correlation of m^6^A modification with either steady state transcript levels or RNA stability. Altogether, these results suggest that m^6^A mediates translational control. Moreover, as our miCLIP maps were generated from highly dT-selected RNAs, we infer that this may reflect modifications that are mostly present in cytoplasmic transcripts available for binding to YTHDF.

### *Drosophila* m^6^A mediates optimal translational activation

The fact that the majority of *Drosophila* m^6^A is present in 5’ UTRs, in contrast to mammalian m^6^A which is predominantly in 3’ UTRs, is suggestive of its role in influencing translation. However, the above genomic analyses are correlational in nature, and do not directly connect m^6^A to gene regulation. One scenario is that m^6^A, being enriched amongst poorly translated mRNAs, is a suppressive mark. However, an alternative regulatory logic is that m^6^A is a positive modification that is preferentially deposited on transcripts with lower translational efficiency, thus making the potential impact of translational enhancement more overt.

We first evaluated whether m^6^A might exert a global impact on translation. We exploited our *mettl3-KO* S2-S cells and monitored newly synthesized proteins using puromycin incorporation (Schmidt et al., 2009). Interestingly, we observe a clear difference between bulk steady-state protein accumulation and nascent protein synthesis between these wild-type and m^6^A-mutant cells. In particular, when analyzing similar amounts of total cellular protein, we observed that *mettl3-KO* cells consistently generated less than half the amount of newly synthesized proteins at different puromycin concentrations (**Figure 6H-I**). Western blotting for tubulin, a non-m^6^A target, verified similar steady-state accumulation between wild-type and knockout cells (although slightly less protein accumulated during puromycin incubation as expected), while Mettl3 blotting confirmed knockout cell status. These data suggested that m^6^A may enhance translation, even though it predominates on genes with poorer translational capacity.

### *Drosophila* YTHDF enhances output of an m^6^A reporter in a YTH-dependent manner

To test if YTHDF might be an effector of m^6^A-mediated translational regulation, we implemented a transgenic assay. We used a reporter backbone consisting of GFP under control of the tubulin promoter (*tub-GFP*), a transgene that is broadly and relatively evenly expressed in the animal (Brennecke et al., 2005). In this genetic background, we can coexpress factors in a spatially defined subpattern, to assess regulatory impact on the transgene. When we stain wing imaginal discs bearing a naive reporter, and expressing UAS-YTHDF in the dorsal compartment (using *ap-Gal4*) or along the anterior-posterior boundary (using *ptc-Gal4*), we do not observe substantially different GFP protein accumulation in cells co-expressing wild-type or mutant YTHDF, compared to non-Gal4 cells as internal control territories (**Figure 7A**), or the documented reporter pattern by itself (Brennecke et al., 2005; Lai et al., 2005).

**Figure 7.**
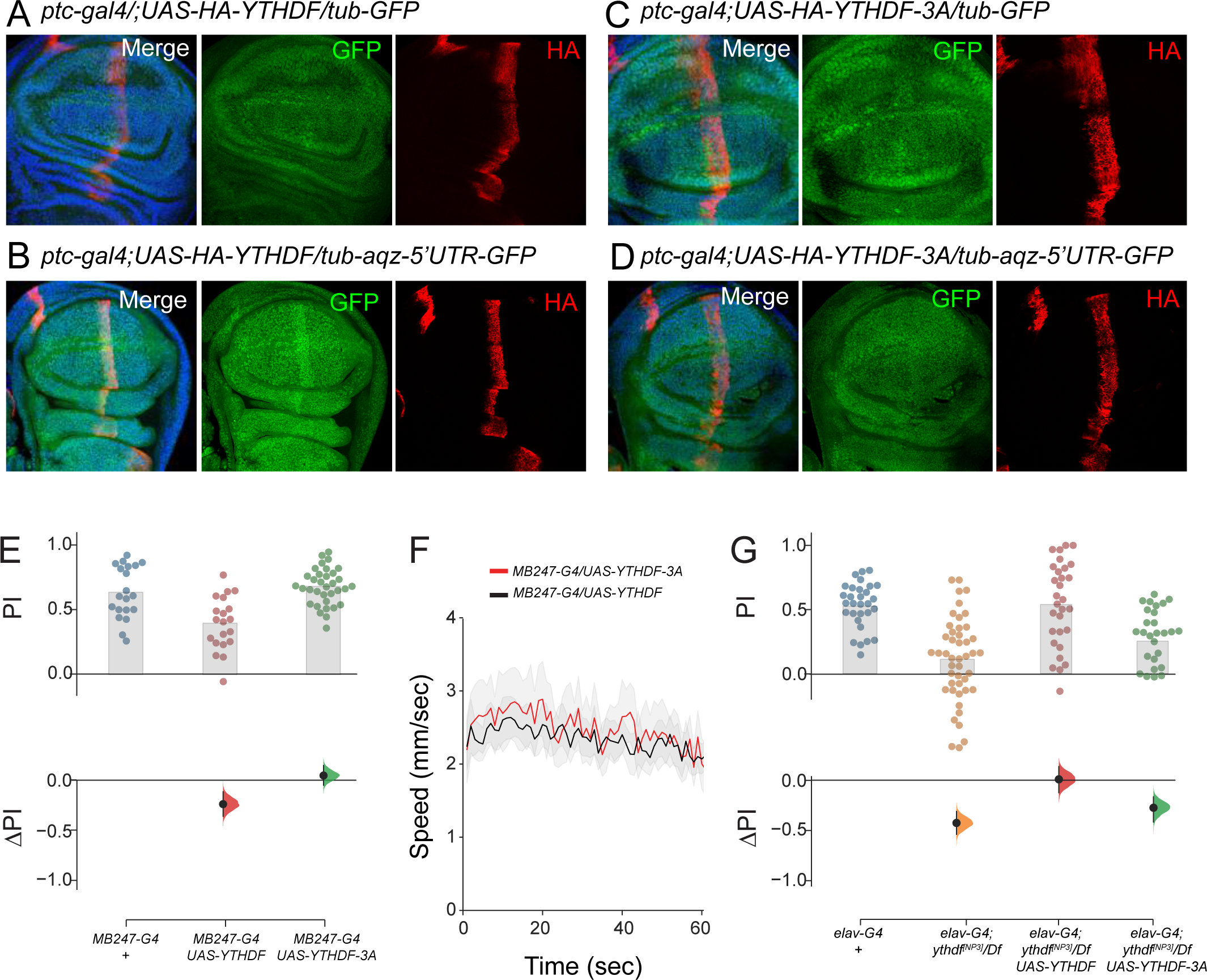
m^6^A binding by YTHDF mediates translational enhancement and STM (A-D) Wing imaginal discs expressing tub-GFP (GFP stained in green), *ptc-Gal4>UAS-HA-YTHDF* wild type or *3A* mutants (HA stained red) and DAPI (blue); non-YTHDF-expressing territories serve as internal controls. (A-B) Expression of wild-type YTHDF has a marginal effect on the parental *tub-GFP* reporter (A), but enhances the *tub-aqz-5’UTR* reporter (B). (C-D) The m^6^A-binding mutant YTHDF-3A has little effect on either reporter. (E) Specific expression of YTHDF, but not YTHDF-3A, in the mushroom body impairs STM in 20d flies. (F) Expression of YTHDF-wt/3A proteins do not affect locomotion behavior. (G) Rescue assay using *elav-Gal4*. Pan-neuronal expression of wild-type YTHDF, but not YTHDF-3A, rescued STM defects in *ythdf* hemizygote mutants at 20 days.

We inserted the 5’ UTR of *aaquetzalli (aqz)*, a validated YTHDF target that exhibited high enrichment of m^6^A (**Figure 5**), into the *tub-GFP* reporter. Aqz is required for cell polarity and neural development (Mendoza-Ortiz et al., 2018). The *tub-aqz-5’UTR-GFP* transgene expressed GFP broadly and the levels were not noticeably different from the parent transgene. However, when we introduced these into *ptc>YTHDF* background, GFP was elevated specifically within the YTHDF-expressing domain (**Figure 7B**). This was consistent with a role for YTHDF in translational enhancement of this m^6^A target.

To test if this was due to specific activity of YTHDF, we generated a transgene containing the three YTH pocket mutations, which we showed abrogates association to m^6^A *in vitro* (**Figure 1C**) and to validate m^6^A-bearing transcripts in cells (**Figure 5C**). YTHDF-3A protein accumulated to a similar level as wild type, and was also similarly neutral as its wild-type counterpart when tested on the parent *tub-GFP* reporter (**Figure 7C**). However, mutant YTHDF- 3A was unable to enhance GFP protein output from the *tub-aqz-5’UTR-GFP* transgenic reporter (**Figure 7D**). These data support the notion that YTHDF recognizes m^6^A-bearing 5’UTR targets for translational enhancement.

### An autonomous, m^6^A-dependent, neural function for YTHDF in memory

The availability of companion wild-type and mutant YTHDF transgenes allowed us to conduct further genetic tests of the connection between m^6^A readout and memory.

Overexpression of either YTHDF transgene, when expressed ubiquitously (with *da-Gal4*) or with tissue-specific drivers (e.g. *ptc-Gal4*, *ap-Gal4*), did not substantially impair viability or developmental patterning; and pan-neuronal expression of YTHDF using *elav-Gal4* did not affect lifespan (**Supplementary Figure 6**). Therefore, even though we could detect a selective m^6^A-dependent impact of YTHDF on reporter transgenes, the overall impact of elevated YTHDF expression is not sufficiently severe as to interfere with normal developmental programs, or is otherwise within the range of developmental compensation. This mirrors the lack of substantial consequences of removing YTHDF.

Bearing in mind that YTHDF-mediated regulation has particular impact on memory, we evaluated these transgenes for effects on this behavior. Strikingly, *mb247-Gal4>YTHDF* flies exhibited compromised memory formation at 20 days, while *mb247-Gal4>YTHDF-3A* flies were normal (**Figure 7E**). This behavioral defect appears to be specific: flies overexpressing wild-type and mutant YTHDF exhibited similar locomotor activity when quantified over 60 s of tracking (**Figure 7F**). Thus, ectopic YTHDF disrupts memory in an m^6^A-dependent manner.

We extended these assays by performing cell-type specific transgenic rescue assays. Building on our observation that neural-knockdown of YTHDF phenocopied the STM defects seen in whole-animal mutations (**Figure 3G**), we introduced *elav-Gal4* and *UAS-YTHDF-wt* or *UAS-YTHDF-3A* transgenes into *ythdf* hemizygous nulls. The *ythdf* nulls carrying *elav-Gal4* had defective memory, indicating the Gal4 transgene does not improve this behavioral output; this was important to rule out, since at least some other *Drosophila* neuronal phenotypes are modified by Gal4 alone (Bollepalli et al., 2017; Kramer and Staveley, 2003). With this control background as reference, we found that the STM deficits of *ythdf* nulls could be rescued by pan-neuronal restoration of YTHDF, restoring normal memory (**Figure 7G**). In contrast, *elav>YTHDF-3A* transgenes did not restore normal STM capacity to *ythdf* mutants (**Figure 7G**).

These results indicate that m^6^A binding is critical for YTHDF function in memory-storing cells.

Taken together, these genetic and genomic data provide evidence that the m^6^A/YTHDF pathway mediates translational enhancement in *Drosophila*, and that normal levels of this system are of particular importance in neurons to maintain normal memory.

## Discussion

### Distinct local contexts, genic location, and regulatory impact for m^6^A in different metazoans

Despite tremendous interests in the regulatory utilities and biological impacts of mRNA methylation, there has been relatively little study from invertebrate models. Given that the m^6^A pathway seems to have been lost from *C. elegans*, *Drosophila* is an ideal choice for this. Since the initial report that *mettl3* mutants affect germline development (Hongay and Orr-Weaver, 2011), we and others showed that *Drosophila* harbors an m^6^A pathway similar to that in mammals, but simplified in that it has a single nuclear and cytoplasmic YTH reader (Haussmann et al., 2016; Kan et al., 2017; Lence et al., 2016). Nevertheless, *Drosophila* has proven to be a useful system to discover and characterize novel m^6^A factors (Guo et al., 2018; Kan et al., 2017; Knuckles et al., 2018; Lence et al., 2016). Expanding the breadth of model systems can increase our appreciation for the utilization and impact of this regulatory modification.

It is widely presumed, based on mammalian profiling, that metazoan m^6^A is enriched at stop codons and 3’ UTRs. However, our new high-resolution maps indicate that 5’ UTRs are by far the dominant location of methylation in mature *Drosophila* mRNAs. Although further study is required, many of these m^6^A 5’ UTR regions coincide with our previous embryo miCLIP data (e.g. Supplementary Figure 9), while other miCLIP CIMs calls located in other transcript regions (Kan et al., 2017) proved usually not to be Mettl3-dependent. Thus, our data indicate a fundamentally different distribution of m^6^A in *Drosophila* mRNAs compared to mammals.

While mammalian m^6^A clearly elicits a diversity of regulatory consequences, depending on genic and cellular context and other factors, a dominant role is to induce target decay through one or more cytoplasmic YTH readers. This harkens back to classic observations that m^6^A is correlated with preferential transcript decay, and more recent data that loss of m^6^A writers or cytoplasmic YTH readers results in directional upregulation of m^6^A targets. However, several lines of study did not yield convincing evidence for a broad role for the *Drosophila* m^6^A pathway in target decay. Instead, the dominant localization of m^6^A in fly 5’ UTRs is suggestive of a possible impact in translational regulation. Our genomic and genetic evidence support the notion that m^6^A is preferentially deposited in transcripts with overall lower translational efficiency, but that m^6^A/YTHDF potentiate translation. However, we can rationalize a regulatory basis for these apparently opposite trends, if the greater modulatory window of poorly translated loci is utilized for preferred targeting by m^6^A/YTHDF.

As is generally the case for mammalian m^6^A, the choice of how appropriate targets are selected for modification, and which gene regions are preferentially methylated, remains to be understood. The minimal context for m^6^A is insufficient to explain targeting, and as mentioned also seems to be different between *Drosophila* and vertebrates. A further challenge for the future will be to elucidate a mechanism for m^6^A/YTHDF-mediated translational regulation. This will reveal possible similarities or distinctions with the multiple strategies proposed for translational regulation by mammalian m^6^A, which include both cap-independent translation via 5’ UTRs during the heat-shock response via eIF3 (Meyer et al., 2015) or YTHDF2 (Zhou et al., 2015); cap-dependent mRNA circularization via Mettl3-eIF3H (Choe et al., 2018); and activity-dependent translational activation in neurons (Shi et al., 2018).

### Roles for the m^6^A/DF1 pathway in learning and memory

Recent studies have highlighted neuronal functions of mammalian m^6^A pathway factors (Du et al., 2019; Widagdo and Anggono, 2018). There is a growing appreciation that mouse mutants of multiple components in the m^6^A RNA-modification machinery affect learning and memory (Koranda et al., 2018; Shi et al., 2018; Walters et al., 2017; Widagdo et al., 2016; Zhang et al., 2018c). Here, we provide substantial evidence that, in *Drosophila*, neural m^6^A is critical for short-term memory (STM). We specifically focused on STM as this paradigm has been extensively characterized in *Drosophila*. Mouse studies have almost exclusively examined effects on LTM, and these two memory phases are mechanistically distinct (Isabel et al., 2004; Quinn and Dudai, 1976). One main distinction is that LTM requires protein synthesis after training, while STM does not. So, while direct comparisons between the two systems are not possible, it is nevertheless instructive to consider the parallels and distinctions of how m^6^A facilitates normal memory function in these species. This is especially relevant given that both mouse and fly central nervous systems require a cytoplasmic YTH factor for memory.

In mice, the m^6^A writer Mettl3 is reported to enhance long term memory consolidation, potentially by promoting the expression of genes such as *Arc*, *c-Fos* and others (Zhang et al., 2018c). Another study found that Mettl14 is required for LTM formation and neuronal excitability (Koranda et al., 2018). Conversely, knockdown of the m^6^A demethylase FTO in the mouse prefrontal cortex resulted in enhanced memory consolidation (Widagdo et al., 2016). Amongst YTH m^6^A readers, YTHDF1 was shown to induce the translation of m^6^A-marked mRNA specifically in stimulated neurons (Shi et al., 2018). In cultured hippocampal neurons, levels of YTHDF1 in the PSD fraction were found to increase by ∼30% following KCl treatment. This suggests that YTHDF1 concentration at the synapse could be critical for regulating the expression levels of proteins (such as Camk2a) that play roles in synaptic plasticity (Giese and Mizuno, 2013). Taken together, these studies suggest that the m^6^A modification pathway is a crucial transcriptomic mechanism of LTM consolidation that optimizes animal behavioral responses.

Of note, the genetics possible in *Drosophila* permit both comprehensive and stringent analyses. Larger sample sizes in *Drosophila* allow markedly improved statistical power (Button et al., 2013). Thus, in our study, we systematically analyze all writer and reader factors, and reveal a notable functional segregation that the cytoplasmic reader YTHDF seems to be major effector of Mettl3/Mettl14 m^6^A in memory. Given that *ythdf* mutants otherwise exhibit few overt developmental or behavioral defects in normal or sensitized backgrounds (while *ythdc* mutants generally phenocopy *mettl3/mettl14* mutants) its role in memory is a novel, surprising insight into the contribution of YTHDF to a critical adaptive function. Moreover, we can pinpoint the spatial requirements of m^6^A for memory, by showing that (1) neuronal restoration of Mettl3 or YTHDF to their respective whole-animal knockouts restores normal STM, and (2) neuronal-specific and mushroom body-specific depletion of *mettl3/ythdf* can induce defective STM. Finally, our findings that both loss and gain of function manipulations in the mushroom body are sufficient to impair *Drosophila* STM points towards a homeostatic role of m6A modulation in *Drosophila* STM.

We observed that learning and memory defects in fly m^6^A mutants are age-dependent, which has not been reported in mammals. Although many physiological capacities decline with life history, these seem to be decoupled from other age-related phenotypes, since mutation of *ythdf* or neural overexpression of YTHDF can interfere with STM but do not substantially impact lifespan or locomotion. Similar for other classical learning and memory genes such as *rutabaga* (Tamura et al., 2003), aging led to a progressive impairment in STM in m^6^A mutants. However, in contrast to other classical learning genes, STM impairment in m^6^A mutants was absent in freshly eclosed flies and only became apparent with progressing age.

One interpretation is that there is a cumulative effect of deregulated m^6^A networks that has an progressive impact specific to mushroom-body neurons. To gain further mechanistic insights, future studies will need to examine age-related changes in gene expression and/or translation, in a cell-specific manner. It remains to be seen whether specific deregulated targets downstream of YTHDF have large individual effects, or whether the memory deficits arise from myriad small effects on translation. YTHDF-CLIP and ribosome profiling from the CNS may prove useful to decipher this. Assuming that loss of translational enhancement of m^6^A/YTHDF targets mediates STM defects, one possibility, to be explored in future studies, is that some targets may already be known from prior genetic studies of memory (Tumkaya et al., 2018).

## Materials and Methods

### m^6^A reader constructs

We obtained full-length cDNAs obtained by PCR from a cDNA library for YTHDC (encoding YTHDC1-PA, 721aa), and YTHDF (encoding YTHDF-PA, 700aa) into pENTR vector. We then used site-directed mutagenesis (primers listed in **Supplementary Table 6**) to generate pENTR-YTHDC-3A (w276A w327A L338A), and pENTR-YTHDF-3A (w404A-W459A-w464A).

These were transferred into the *Drosophila* Gateway vector pAGW (N-terminal GFP fusion) and pAHW (N-terminal HA tag) to make all combinations of tagged wild-type and “3A” mutant versions for expression in *Drosophila* cell culture. We also cloned the YTHDF sequences into pTHW (N-terminal HA fusion) to generate the UAS-HA-YTHDF and UAS-HA-YTHDF-3a for transgenic expression. For mammalian expression, we cloned wild-type and “3A” mutant versions of YTHDC and YTHDF into pcDNA5/FRT/TO with an N-terminal 3xFlag tag.

### m^6^A probe pulldown with fly YTHDF and YTHDC

For initial interaction tests of wild-type Flag-YTHDC/DF with A/m^6^A RNA probes, we seeded 4 million HEK293T cells in a 10 cm dish 24 h prior to calcium phosphate transfection, and used 10 µg of plasmid DNA (pcDNA-3xFlag-YTHDC/DF constructs) per 10 cm dish. Cells were harvested 24 h post transfection. Cells from one 10 cm dish and 0.6 mL of lysis buffer were used per condition. Cells were lysed in NP-40 lysis buffer (50 mM Tris-HCl pH 7.5, 150 mM NaCl, 0.5 % NP-40, 5 mM MgCl_2_, EDTA-free protease inhibitor tablet (Roche), 1 mM PMSF) on ice. Clarified lysate was then incubated with A or m^6^A containing RNA probe (**Supplementary Table 6**) on ice at 1 µM concentration for 20 min. Reactions were then irradiated with 365 nm UV (Spectroline ML-3500S) on ice for 10 min. The reaction was then incubated with 60 µL of high capacity streptavidin agarose 50 % bead slurry (Pierce #20357) at 4°C on a rotatory wheel for 3 h. The beads were then washed with 1% SDS in TBS (3x 1 mL), 6 M urea in TBS (3x 1 mL), and TBS (3x 1 mL). The RNA-bound proteins were eluted by boiling the beads in 50 µL of 1x Laemmli sample buffer (80 mM Tris-HCl pH 6.8, 2 % SDS, 10 % glycerol, 5 % B-mercaptoethanol, 0.02 % bromophenol blue) at 95 °C for 5 min. The input, flow-through, and eluates were separated by SDS-PAGE and analyzed by Western blotting using mouse α-FLAG M2 (Sigma #F1804).

In the experiment where 3A mutants of YTHDF and YTHDC were tested, cells from two 10 cm dishes and 1.2 mL of lysis buffer were used per condition. The cross-linking and pull-down were performed as described above. AAACU and AA-m^6^A-CU sequences were used for this experiment.

### Proteomic profiling of the fly m^6^A interactome

*Mass spectrometry analysis*. For proteomics experiments with *Drosophila* S2 cells, we adapted our previously described method (Arguello et al., 2017). S2 cells were lysed by cryomilling. The resulting cell powder (750 mg) was first extracted with 1.5 mL of low-salt extraction buffer (20 mM Tris-HCl pH 7.5, 10 mM NaCl, 2 mM MgCl-_2_, 0.5 % Triton X-100, 10 % glycerol, protease inhibitor tablet (Roche) and phosphatase inhibitor (Pierce)), and then 1 mL of high-salt extraction buffer (50 mM Tris-HCl pH 7.5, 420 mM NaCl, 2 mM MgCl-_2_, 0.5 % Triton X-100, 10 % glycerol, protease inhibitor tablet (Sigma) and phosphatase inhibitor (Pierce)). Low-salt and High-salt extracts were pooled, and protein concentration was determined by Bradford assay. The pooled extract was diluted to 3 mg/mL if needed before proceeding to photo-crosslinking.

AAACU or AA-m^6^A-CU oligo probe was added to 2 mL of extract to a final concentration of 1 µM. The reactions were incubated on ice for 20 min prior to photo-cross-linking. The reactions were then irradiated with 365 nm UV (Spectroline ML-3500S) on ice for 15 min. The reaction was then incubated with 60 µL of high capacity streptavidin agarose 50 % bead slurry (Pierce #20357) at 4 °C on a rotatory wheel for 3 h. The beads were then washed with 1% SDS in TBS (3x 1 mL), 6 M urea in TBS (3x 1 mL), and TBS (3x 1 mL). The RNA-bound proteins were eluted with RNase cocktail (Thermo Fisher) in RNase elution buffer (10 mM Tris-HCl pH 7.5, 40 mM NaCl, 1 mM MgCl_2_) at 37 °C for 30 min with periodic agitation.

The proteomics files were searched against *Drosophila melanogaster* database downloaded from UniProt (https://www.uniprot.org/). To plot the mass spectrometry data (Figure 1D), we first removed 242 proteins that were not consistently recovered in both replicate datasets, leaving 353 proteins. To calculate enrichment ratios for proteins identified by only one probe, we added 1 to all spectral count values. Proteins that do not exhibit differential binding to the A/m^6^A probes cluster around the plot origin, and we thresholded at 2.0× the interquartile range.

### Analysis of m^6^A by liquid chromatography-coupled mass spectrometry

Total RNA was extracted from *Drosophila* S2 cells and whole female fly (one-week old) using TRIzol reagent and subjected to DNase treatment (Thermo Fisher #AM1907). The mRNA was then isolated through two rounds of poly-A selection using the oligo-d(T)_25_ beads (NEB #S1419S). The RNA was digested with nuclease P1 (Wako USA #145-08221) and dephosphorylated with Antarctic phosphatase (NEB #M0289S). Briefly, 1 µg of RNA was digested with 2 units of nuclease P1 in buffer containing 7 mM NaOAc pH 5.2, 0.4 mM ZnCl_2_ in a total volume of 30 µL at 37°C for 2 h. 3.5 µL of 10x Antarctic phosphatase buffer and 1.5 µL of Antarctic phosphatase was then directly added to the reaction and incubated at 37°C for another 2 h.

Quantitative LC-MS analysis of m^6^A was performed on an Agilent 1260 Infinity II HPLC coupled to an Agilent 6470 triple quadrupole mass spectrometer in positive ion mode using dynamic multiple reaction monitoring (DMRM). The ribonucleosides in the digested RNA samples were separated by a Hypersil GOLD™ C18 Selectivity HPLC Column (Thermo Fisher #25003-152130; 3 µm particle size, 175 Å pore size, 2.1 x 150 mm; 36 °C) at 0.4 mL/min using a solvent system consisting of 0.1 % formic acid in H_2_O (A) and acetonitrile (B) based upon literature precedent (Su et al., 2014). The operating parameters for the mass spectrometer were as follows: gas temperature 325 °C; gas flow 12 L/min; nebulizer 20 psi and capillary voltage 2500 V, with fragmentor voltage and collision energy optimized for each different nucleoside.

The nucleosides were identified based on the transition of the parent ion to the deglycosylated base ion: m/z 282 → 150 for m^6^A and m/z 268 → 136 for A. Calibration curves were constructed for each nucleoside using standards prepared from commercially available ribonucleosides. The level of m^6^A was determined by normalizing m^6^A concentration to A concentration in the sample.

### Co-immunoprecipitation tests

We generated pAC5.1-Fmr1-3xflag-3xHA, FMR full-length sequence with C-terminal 3xFlag/3XHA was obtained by PCR from pNIK1147 (20XUAS-FMR1-3XFLAG-3XHA, gift of Nicholas Sokol) (Luhur et al., 2017) into pAC5.1 with KpnI/NotI. To generate pAc5.1A-GFP-Fmr1, we used Gibson assembly to assemble GFP and FMR full-length sequence into pAc5.1A. To test YTHDF and FMR1 association, we transfected S2 cells in each well of a 6-well plate with 1 µg each of HA/GFP-tagged constructs using Effectene (Qiagen). After incubation for 3 days, cells were washed with PBS and lysed with co-IP lysis buffer (30mM HEPES, pH 7.5, 150mM KOAc, 2mM Mg(OAc)2, 5mM DTT, 0.1% NP40, 1X protease inhibitor (Roche) on ice for 30 min, and then followed by two centrifugation at 20,000 g for 10 min at 4 °C. The cleared cell lysates are incubated with mouse α-HA (F-7, Santa Cruz, #Sc-7392), or mouse α-GFP (3E6, Invitrogen, A11120) conjugated beads for 2 h. Beads were washed with co-IP lysis buffer for five times and separated the sample into 90% for RNA, 10% for WB in the last wash. Beads for WB was resuspended in 40 ul 1XSDS sample buffer and incubated at 95°C for 5 min. For Western blotting, we used mouse α-HA-Peroxidase (1:5000, H6533-1VL, Sigma), rabbit α-GFP (1:1000, A11122, Invitrogen), mouse α-Fmr1 (6A15,1:1000, Santa Cruz, sc-57005). Goat α-rabbit and α-mouse secondary antibodies conjugated to HRP (Jackson) were used at 1:5000.

### Drosophila stocks

*YTHDF[NP3], FRT40A* and *mettl14[sk1]/Tb-RFP cyo,w+* were previously described (Kan et al., 2017). *YTHDC[ΔN]/Dfd-YFP, TM3*; *mettl3[Δcat]/TM6C*, *mettl3[null]/TM6C*, *mettl14[fs]/Tb-RFP cyo,w+*; *UAS-mettl3-HA* were a gift of Jean-Yves Roignant (Lence et al., 2016); UAS-YTHDC:2XHA was a gift of Matthias Soller (Haussmann et al., 2016). All of these were genotyped in trans to deficiencies (see **Supplementary Table 6**) to confirm the absence of the wild-type allele. Other stocks were obtained from the Bloomington *Drosophila* Stock Center.

Deficiency lines: *Df(3R)Exel6197* (BL-41590, removes *mettl3*), *Df(3L)ED208* (BL-34627, removes *ythdc*), *Df(3R)BSC461* (BL-24965, removes *ythdf*) and *Df(3R)BSC655* (BL-26507, removes *ythdf*). TRiP knockdown lines: *mettl3* (BL-41590), *mettl14* (BL-64547), *YTHDC1* (BL-34627) and *YTHDF* (BL-55151). Gal4 lines: *elav-gal4* (BL-8765), *elav[C155]-GAL4* (BL-458), *ptc-GAL4* (BL-2017), *tub-Gal4* (BL-5138), *da-Gal4*l(BL-55851) and *ap-gal4* (BL-3041).

To generate the *tub-aqz-GFP* reporter, we cloned the *aqz* 5’ UTR (chr3R:8,818,731-8,820,682) into the 5’UTR position of the tub-GFP vector (KpnI/BamHI). tub-aqz-GFP, UAS-HA-YTHDF and UAS-HA-YTHDF-3A (described above) were injected into *w[1118]* with Δ2-3 helper plasmid to obtain transformants (Bestgene, Inc.) Flies were raised on standard cornmeal-based food medium containing 1.25% w/v agar, 10.5 % w/v dextrose, 10.5% w/v maize and 2.1% w/v yeast at 60% relative humidity.

### Survival experiments

All *Drosophila* survival experiments were performed with at least 90 mated female flies per genotype at 23°C. Throughout the lifespan assessment, flies were kept in vials in groups of 10 and transferred to a new food vial every second or third day. The number of surviving flies was counted after each transfer. Average lifespan was calculated using the DABEST estimation statistics package (Ho et al., 2019). The data were plotted to compare the average survival of each tested genotype against the average survival of the *w[1118]* control stock that was assayed in parallel at the same time.

### The multifly olfactory trainer (MOT) conditioning apparatus

The MOT apparatus was designed to allow the monitoring of *Drosophila* behavior throughout olfactory conditioning in a controlled environment (Claridge-Chang et al., 2009). Flies were assayed in conditioning chambers, whereby the arena of each chamber was 50 mm long, 5 mm wide and 1.3 mm high (**Supplementary Figure 3A**). The floor and ceiling of each chamber was composed of a glass slide printed with transparent indium tin oxide electrodes (Walthy, China). Each side of the electrode board was sealed by a gasketed lid that formed a seal around the gap between the electrode board and the chamber wall. Facilitated by carrier air, the odors entered the chamber via two entry pipes and left the chamber through two vents that were located in the middle of the chamber. Up to four MOT chambers were stacked onto a rack which was connected to the odor and electric shock supply (**Supplementary Figure 3B**). Chambers were illuminated from the back by two grids of infrared LEDs. Fly behaviour inside the chambers was recorded with an AVT F-080 Guppy camera (Allied Vision) that was connected to a video acquisition board (PCI-1409, National Instruments). Electric shock during odor presentation is delivered when the animals walk on the electrode contacts. Olfactory preference was measured by tracking the movement of individual flies and scored automatically by using the in-house designed CRITTA tracking programme (Krishnan et al., 2014).

### MOT odor delivery and odor concentrations

Conditioning was done as previously described (Claridge-Chang et al., 2009), with some protocol modifications. The rack with stacked conditioning chambers was connected to an olfactometer that was used to deliver precisely timed odor stimuli (**Supplementary Figure 3B**). The conditioning odors methylcyclohexanol (MCH) and 3-octanol (OCT) were carried by dry, compressed air and routed through mass flow controllers (MFC; Sensirion AG, Sweden). Carrier air flow was controlled with two 2 L/min capacity MFCs and pushed through a humidifying gas washing bottle containing distilled water (Schott Duran) at 0.6 L/min. Odor streams were controlled with 500 mL/min MFCs and pushed through glass vials containing pure liquid odorants (either MCH or OCT respectively). Prior to conditioning, the odor concentrations were adjusted to ensure that flies did not display a strong preference for one of the odors over the other prior to training. Odor administration was carried out with the following MFC settings: OCT left side 25-35 mL/min; OCT right side 30-40 mL/min; MCH left side 50-60 mL/min; MCH right side 50-60 mL/min. Odor presentation at the behavioral chamber arms was switched with computer-controlled solenoid valves (The Lee Company, USA). The MFCs were regulated via CRITTA (LabView software).At *ad hoc* intervals between experiments, odor concentrations were measured with a photoionization detector (PID, RAE systems; PGM-7340). The experiments were performed with a relative concentration of 14–16 parts per million (ppm) for MCH and 6–8 ppm for OCT in the chambers. A relative humidity of 70–75 % was maintained via regulation of the air flow; this was monitored (*ad hoc*, between experiments) by using a custom humidity sensor with a custom LabVIEW code (National Instruments, US).

### Classical olfactory conditioning in the multifly olfactory trainer and data visualization

Classical olfactory conditioning has been described previously (Claridge-Chang et al., 2009; Tully and Quinn, 1985; Tumkaya et al., 2018). Before each experiment, flies were briefly anesthetized on ice and six flies were loaded into each conditioning chamber (**Supplementary Figure 3C**). Each conditioning experiment began with an acclimatization (baseline test) phase where *Drosophila* were exposed to both odors in the absence of a shock stimulus.

Subsequently, in the first stage of training the chambers were flushed with carrier air and flies were exposed to either MCH or OCT in the presence of a shock stimulus (12 shocks at 60 V during a 60 sec time interval). During the second stage of training the shock odor was removed through carrier air and the flies were exposed to the other odor in the absence of a shock stimulus. After removal of the odor and air-puff agitation, flies were tested for shocked-odor avoidance. The flies were given a choice between the two odors and average shocked-odor avoidance was quantified for the last 30 s of the 2 min-long testing phase. The main stages of the conditioning protocol are summarized in **Figure 3B**. The full conditioning protocol is presented in **Supplementary Figure 3D**. The shocked-odor avoidance of flies for each conditioning trial was expressed as a performance index (PI) (Tully and Quinn, 1985); however, instead of a single endpoint, counting was performed on individual video frames over the final 30 s of the testing period. Each trial produced a half PI against the respective conditioned odor (either MCH or OCT) and two half PI’s from consecutive experiments (with different conditioning odors) were combined to a full PI (full PI = half PI OCT + half PI MCH). For data visualization, the distribution of full PI’s was plotted with a 95% CI error presenting a ΔPI between control and test genotypes by using the DABEST estimation-statistics package (Ho et al., 2019).

### *Drosophila* immunostaining

We performed immunostaining as previously described (Lai and Rubin, 2001), by fixing dissected tissues in PBS containing 4% formaldehyde and incubating with the following primary antibodies: mouse α-HA (1:1000, Santa Cruz), and guinea pig α-Mettl3 (1:2000, gift of Cintia Hongay, Clarkson University). Alexa Fluor-488, and-568 secondary antibodies were from Molecular Probes and used at 1:1000. Tissues were mounted in Vectashield mounting buffer with DAPI (Vector Laboratories). Images were captured with a Leica SP5 confocal microscope; endogenous GFP signals were monitored.

### m^6^A individual-nucleotide-resolution cross-linking and immunoprecipitation (miCLIP)

miCLIP libraries were prepared by subjecting RNA samples to the established protocol (Grozhik et al., 2017) with the minor changes described below. Briefly, total RNA was collected from <1 week old *w1118* (wild type) and *mettl3[null]* (mutant) female heads using TRIzol RNA extraction. Poly(A)+ RNA was enriched using two rounds of selection. RNAs were fragmented, incubated with the α-m^6^A (202 003 Synaptic Systems) and crosslinked twice in a Stratalinker 2400 (Stratagene) using 150 mJ per cm^2^. Crosslinked RNAs were immunoprecipitated using Protein A/G magnetic beads (Thermo) and washed under high salt conditions to reduce non-specific binding. Samples were radiolabeled with T4 PNK (NEB), ligated to a 3’ adaptor using T4 RNA Ligase I (NEB), and purified using SDS-polyacrylamide gel electrophoresis (SDS-PAGE) and nitrocellulose membrane transfer. RNA fragments containing crosslinked antibody peptides were recovered from the membrane using proteinase K (Invitrogen) digestion.

Recovered fragments were subjected to library preparation. First-strand cDNA synthesis was performed using SuperScript III (Life Technologies) and iCLIP-barcoded primers, which contain complementarity to the 3’ adaptor on the RNA. cDNAs were purified using denaturing PAGE purification, circularized using CircLigase II (EpiCentre), annealed to the iCLIP Cut Oligo, and digested using BamHI (Thermo). To generate libraries for sequencing, the resulting linear cDNAs were amplified using Accuprime SuperMix I (Invitrogen) and P5 and P3 Solexa primers, and purified using Agencourt AMPure XP beads (Beckman Coulter).

For input libraries, poly(A)+ RNAs were fragmented and directly subjected to radiolabelling and 3’ adaptor ligation. All subsequent steps are as listed above. Libraries were paired-end sequenced on an Illumina HiSeq2500 instrument at the New York Genome Center (NYGC).

### miCLIP bioinformatic analyses

Read processing, mutation calling and annotation of CIMs was performed as described (Grozhik et al., 2017). Briefly, to prepare libraries for mapping, adapters and low quality reads were trimmed using flexbar v2.5. Next, the FASTQ files were de-multiplexed using the pyBarcodeFilter.py script from the pyCRAC suite. Random barcodes were removed from sequencing reads and appended to sequence IDs using an awk script and PCR duplicates were removed using the pyCRAC pyDuplicateRemover.py script. Paired end reads were merged and mapped to the *Drosophila* reference genome sequence (BDGP Release 6/dm6) using Novoalign (Novocraft) with parameters –t 85 and -l 16.

Mutations were called using the CIMS software package (Zhang and Darnell, 2011). To identify putative m^6^A sites, C-to-T transitions with preceding A nucleotides were extracted and filtered such that the number of mutations that support the mismatch (m) > 1 and 0.01 < m/k < 0.5, where k is the number of unique tags that span the mismatch position.

Peaks were called by adapting the Model-based Analysis for ChIP-Seq (MACS) algorithm (Zhang et al., 2008). Mettl3-dependent peaks for head libraries were determined using miCLIP versus input, comparing wild type and *mettl3* libraries and the MACS2 differential binding events program (bdgdiff) with parameters -g 20 and -l 120. Lastly, peaks were split using PeakSplitter (version 1.0, http://www.ebi.ac.uk/research/bertone/software).

To generate nucleotide content plots, filtered C-to-T transitions with preceding A nucleotide (as mentioned above) that mapped within the top 100 or 1000 Mettl3-dependent peaks were chosen to describe the nucleotide content surrounding CIMs. Sequences were obtained using the Drosophila reference genome sequence (dm6) and fed to WebLogo version 2.8.2 with the frequency setting (Crooks et al., 2004).

Custom scripts were used to generate metagene plots. Briefly, to prepare mapped data, each miCLIP bam file was converted to bedGraph format with span of 1 nucleotide. To prepare features, for each gene, the longest transcript model was selected and divided into 5’ UTR, CDS and 3’ UTR segments according to Ensembl transcript models for BDGP6.94. Next, miCLIP read depth mapping to transcripts were selected and scaled such that each 5’ UTR, CDS and 3’ UTR were 200, 1000 and 300 nts. To normalize, the score at each scaled nucleotide was divided by the total score across all 1500 nucleotides. Finally, to yield metagene score across each feature (UTRs and CDS), genes of interest were selected and means were calculated for each nucleotide position. Smoothing functions from the ggplot2 package (Wickham, 2016) were used to visualize metagene analysis.

Pie charts were obtained by mapping peaks to Ensembl transcript models for BDGP 6.94. Since transcript features occasionally overlap, the following order was used to bin peaks into different categories: other (not mapping transcript models), introns, start codons, 5’ UTRs, 3’ UTRs and CDS. Finally, the ggplot2 function geom_bar was used to plot the accounted annotations into a pie chart.

Input normalized miCLIP tracks along with described peaks were used to generate heatmaps using deepTools2 functions computeMatrix and plotHeatmap (Ramirez et al., 2016).

### RNA-seq analysis

Flies of the specified genotypes(*w;;mettl3[null]/+*, *w;;mettl3[null]/mettl3[cat]*, *w;;Df(3R)BSC461/+*, *w;;YTHDF[NP3]/Df(3R)BSC461*, *w;;Df(3L)ED208/+*, *w;;ythdc[ΔN]/Df(3L)ED208*) were aged to one or three weeks at 25°C. Female heads were dissected and collected for total RNA extraction using TRIzol reagent. Sequencing libraries were prepared using the TruSeq Stranded Total RNA Sequencing Kit (Illumina) following the manufacturer’s protocol. Sequencing was performed on a HiSeq 2500 System in paired end read mode, with 100 bases per read at the Integrated Genomics Operation (IGO) at Memorial Sloan Kettering Cancer Center.

RNA sequencing libraries were mapped to the *Drosophila* reference genome sequence (BDGP Release 6/dm6) using HISAT2 (Kim et al., 2015) under the default settings. Gene counts were obtained by assigning and counting reads to the Ensembl transcript models for BDGP6.94 using Rsubread (Liao et al., 2013). Differential gene expression analysis was performed with comparisons as listed in **Supplementary Table 5** using the R package DESeq2 (Love et al., 2014) and applying a strict adjusted p-value cutoff of 0.05.

### CRISPR/Cas9 deletion of *mettl3* in S2-S cells

We used CRISPR/Cas9-mediated mutagenesis as described (Reimao-Pinto et al., 2016) to generate a *mettl3-KO* S2 cell line. Guide RNA sequences are listed in **Supplementary Table 6**. We analyzed 11 candidate clonal lines obtained from subcloning of two initial low-complexity mixed cell populations, and kept the deletion as #3-5 as described in **Supplementary Figure 11**.

### m^6^A-RIP-PCR and RIP-rtPCR

We adapted a protocol from our recent study (Kan et al., 2017). Plasmids of 5 µg were transfected into 6x10[6] S2 cells using Effectene (Qiagen) and incubated for 3 days. Cells were washed with PBS and lysed with IP lysis buffer (30mM HEPES, pH 7.5, 150mM KOAc, 2mM Mg(OAc)_2_, 5mM DTT, 0.1% NP40) supplied with Complete, EDTA-free Protease Inhibitor and 40Uml-1 SUPERase•In RNase Inhibitor (Ambion) on ice for 30 min, followed by 2x10 min centrifugation at 20,000 g at 4^°^C. 10% of the cleared cell lysate were kept as input and the rest was incubated with 15 µl Dynabeads™ Protein G (Thermo Fisher,10004D)(with HA or GFP antibody) for 4 h at 4°C. RNase I (Invitrogen, AM2294) was added to the sample for RNase treatment at 0.2U final concentration. The beads were washed three times using IP lysis buffer and then resuspended in 100 ml lysis buffer. To elute RNA, the beads were mixed with 900 ml of Trizol, vortexed for 1min and incubated at RT for 50 with rotation. RNA extracted and treated were Turbo DNase (Ambion) for 30 min before cDNA synthesis using SuperScript III (Life technology) with random hexamers. PCRs were done using Fusion High-Fidelity Polymerase (ThermoFisher Scientific).

For m^6^A-RIP-qPCR, the mRNAs were immunoprecipitated using α-m^6^A according to the procedure shown above. The IP-mRNAs were then reverse transcribed and amplified following the same protocol. The enrichment of m^6^A was quantified using qPCR as reported. The sequences of qPCR primers are listed in Supplementary Table 6.

### RNA degradation assay

S2 cells were seeded as 3x10^6 cells per well. Actinomycin-D (Gibco, 11-805-017) was added to a final concentration of 5 µM, and cells were collected before or 5 h after adding actinomycin-D. Then the cells were processed as described in ‘RT–qPCR’, except that the data were normalized to the t=0 time point.

### SUnSET assay and Western blotting

For each assay, we incubated 3x10[6] cells in 1ml Schneider’s Medium including 10% FBS for 5 min at 25°C with or without 5 µg/mL puromycin (Gibco™ Sterile Puromycin Dihydrochloride). Cells were then washed twice and incubated for 50 min at 25°C. After washing twice with cold PBS, cells were lysed with 100 µl Lysis buffer (10 mM Tris-HCl, pH 7.5, 300 mM NaCl, 1 mM EDTA, 1% Triton X-100, Protease Inhibitor Cocktail Roche). The cell pellet was resuspended by pipetting and incubated on ice for 30 min, then centrifuged at 16000g for 10 min at 4°C. Protein concentration was measured using Bio-Rad Protein Assay Dye (500-0006) and 2.5 µg proteins were separated on SDS-PAGE and transferred to Immobilon-P membranes. Membranes were blocked for 1 h in TBS containing 5% nonfat milk and 0.1% Tween-20, followed by incubation with mouse α-puromycin (1:1000) overnight at 4°C. Appropriate secondary antibodies conjugated to HRP (Jackson) were used at 1:5000 for 1 h at room temperature, then visualized using chemiluminescence detection (Amersham ECL Prime Western Blotting Detection Reagent). Mouse α-puromycin (2A4, 1:1000, Mouse α-cyclinB (F2F4, 1:100) and mouse α-ß-tubulin (E7, 1:1000) were from DSHB; guinea pig α-Mettl3 (1:5000) was a gift from Cintia Hongay.

## Acknowledgements

We thank the Developmental Studies Hybridoma Bank (DSHB) and Cintia Hongay for antibodies, Jean-Yves Roignant, Matthias Soller and the Bloomington Drosophila Stock Center (BDSC) for fly stocks, Nicholas Sokol for plasmids, Stefan Ameres for S2-S lines and advice on cell mutagenesis. Sara Zaccara, Vladimir Despic and Samie Jaffrey provided helpful advice on generation of miCLIP libraries and discussion about m^6^A. ACC and SO were supported by grants from the Singapore Ministry of Education (MOE2013-T2-2-054 and MOE2017-T2-1-089) and by the Duke-NUS Graduate Medical School. SO was supported by a Khoo Postdoctoral Fellowship Award (Duke-NUS-KPFA/2017/0015). Work in the ECL lab was supported by NIH grants R01-GM083300 and R01-NS083833, and by the MSK Core Grant P30-CA008748.

## Declaration of Interests

The authors declare no competing interests.

## Supplementary Figures and Tables

**Supplementary Figure 1.**
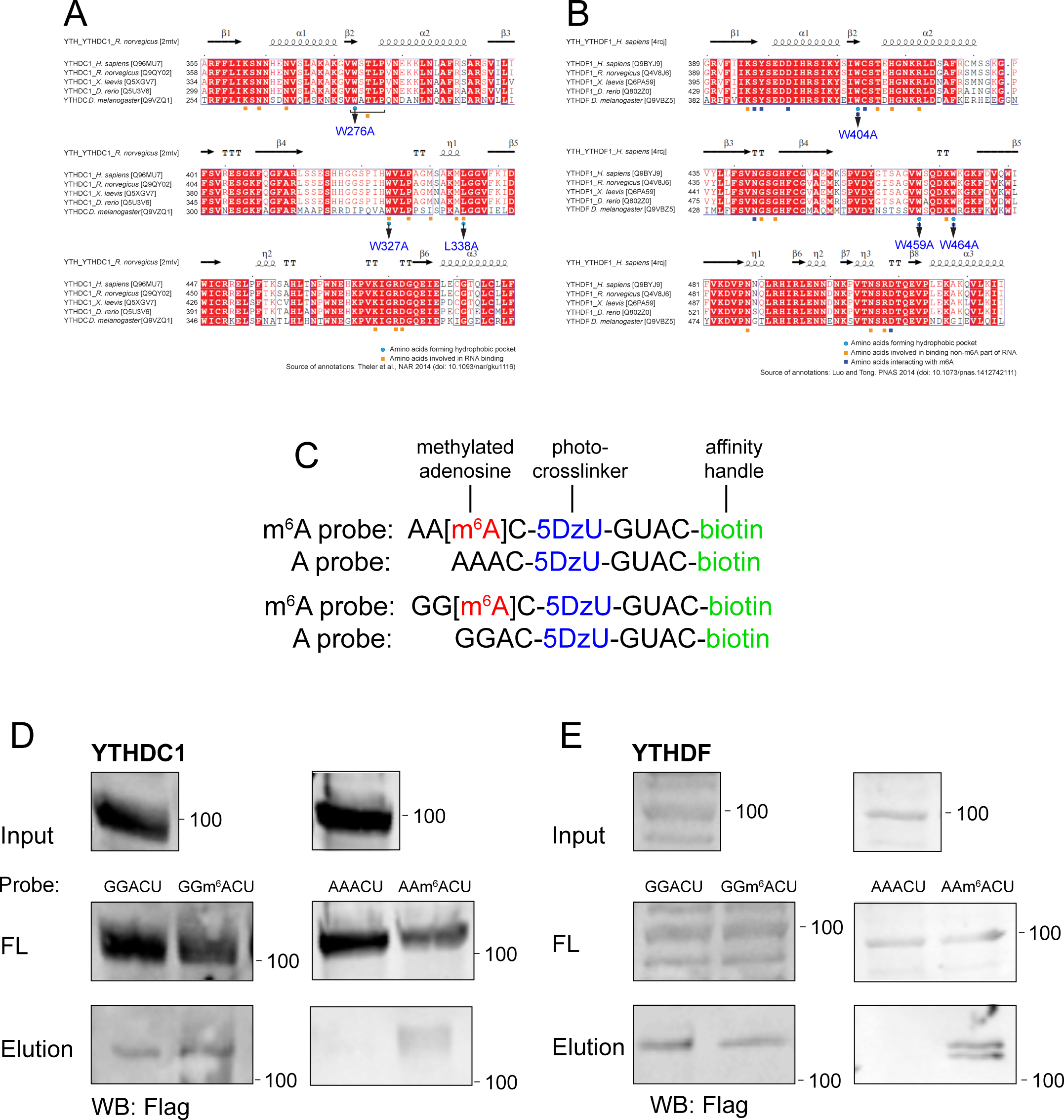
m^6^A binding properties of *Drosophila* YTH proteins. (A-B) YTH domains from nuclear YTHDC (A) and cytoplasmic YTHDF (B) are homologous between *Drosophila* and vertebrates. The critical tryptophan/leucine residues in the YTHDC/YTHDF m^6^A binding pocket were mutated into alanines, as marked with arrows. (C) Sequences of unmodified and modified RNA probes used for photo-crosslinking assays with YTH proteins. (D) YTHDC showed modestly enhanced association to GGm^6^ACU vs. GGACU probes, but showed clearly preferential crosslinking to AAm^6^ACU compared to AAACU probes. (E) YTHDF did not exhibit preferential association to GGm^6^ACU vs. GGACU probes, but was specifically crosslinked to AAm^6^ACU relative to AAACU probes.

**Supplementary Figure 2.**
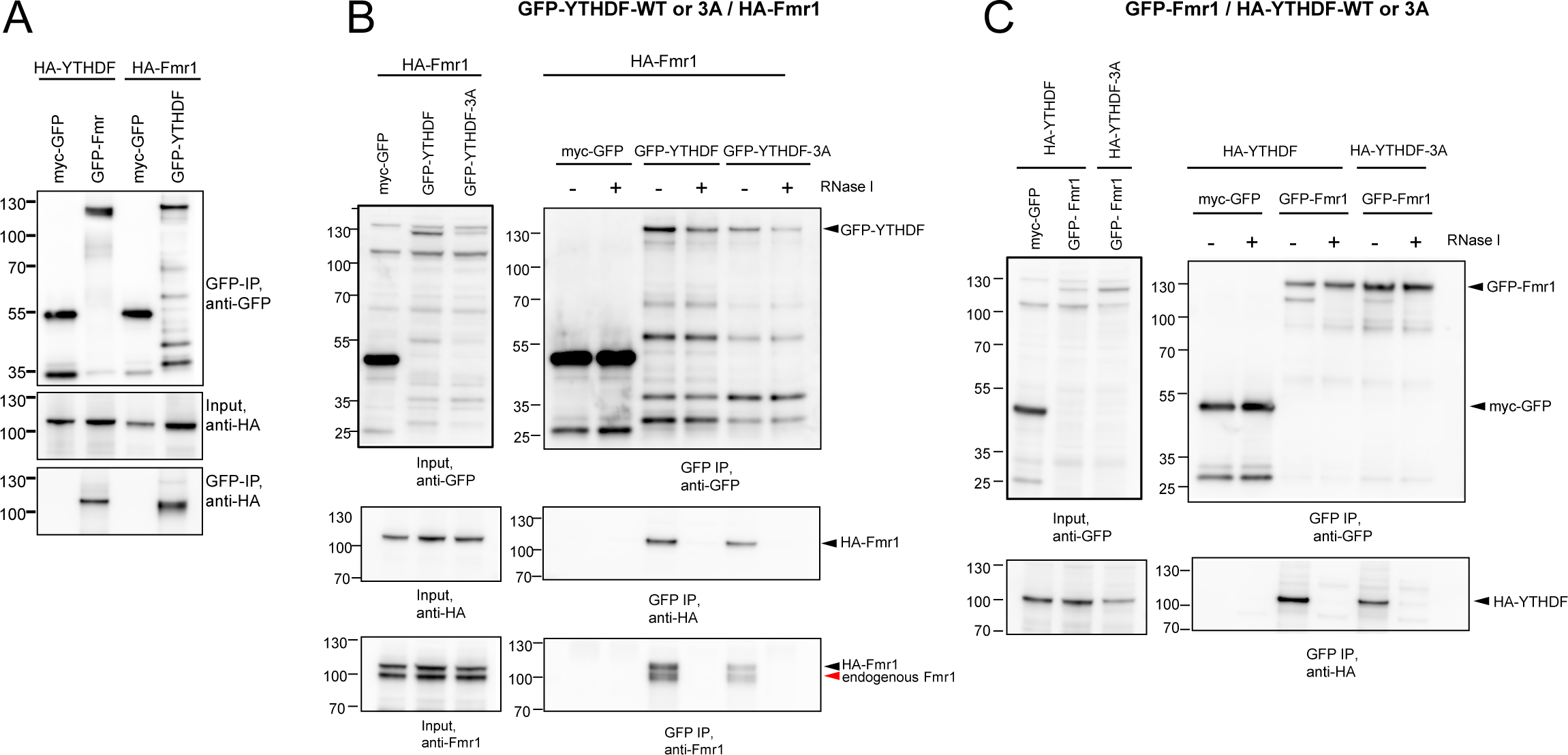
Indirect association of YTHDF and Fmr1. (A) Reciprocal co-immunoprecipitation (co-IP) assays using HA-YTHDF/GFP-Fmr1 and HA-Fmr1/GFP-YTHDF provide evidence that these factors exist in complexes. (B) Co-IP analyses using GFP-YTHDF and mutant 3A forms show that both can be co-IPed with HA-FMR1 and endogenous FMR1. However, these associations are largely abolished by RNase treatment. (C) Similar RNA-dependent associations were detected using GFP-FMR1 and HA-YTHDF and mutant 3A constructs.

**Supplementary Figure 3.**
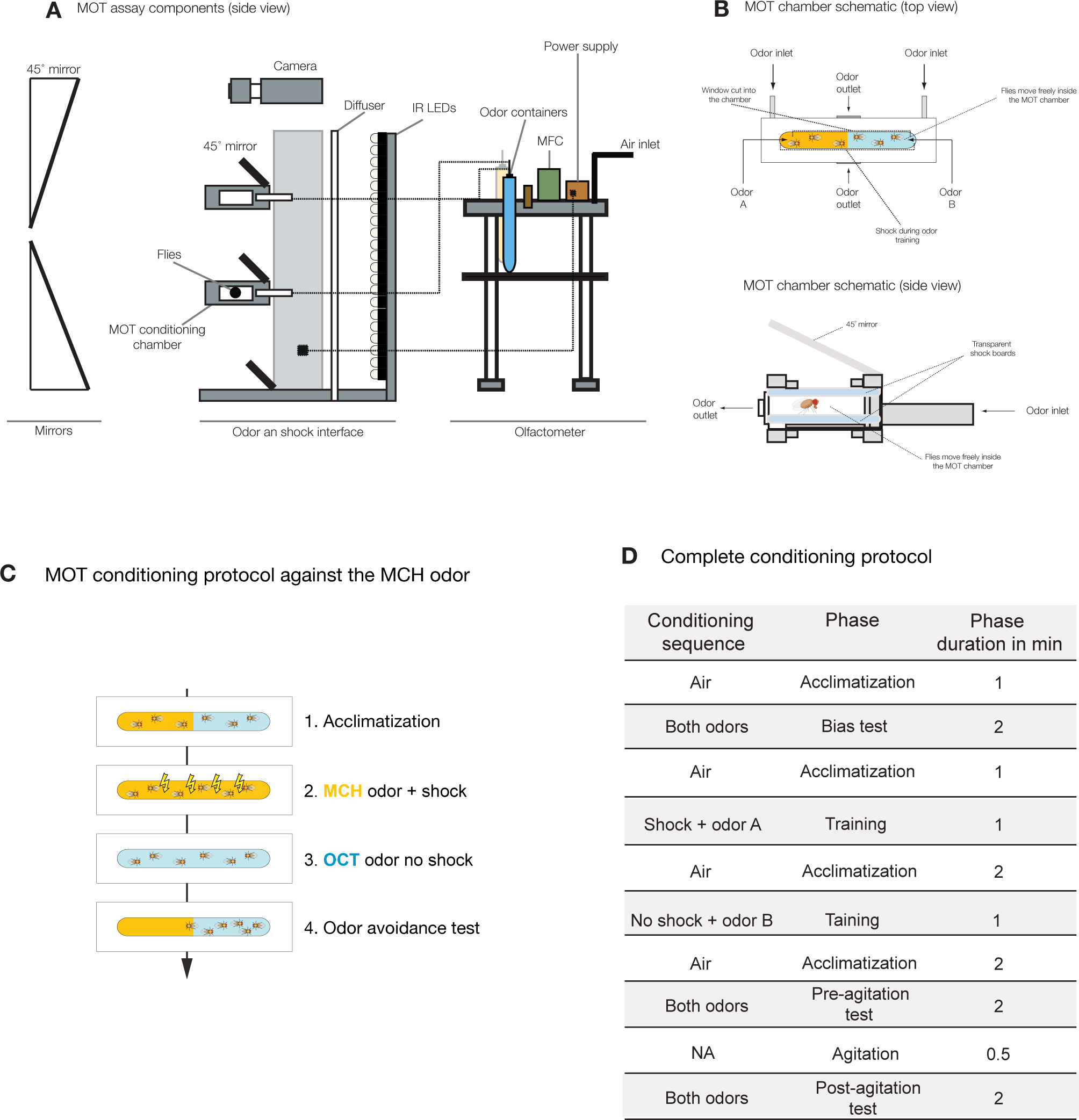
Structure and conditioning protocol of Multifly Olfactory Trainer (MOT) for short term memory evaluation. (A) Overall schematic of MOT assay components. (B) Detailed schematics of the MOT chamber from top and side views. (C) Sequence of a standard MOT conditioning protocol. After *Drosophila* were placed in the chambers and were given 60 s to acclimatize in pure air, one half of the chamber was exposed to MCH while the other half to OCT for during the bias test. After another 90 s of pure air, the whole arena was exposed for 60 s to the conditioned odor in the presence of a foot shock. This was followed by 90 s of air and the exposure of the whole arena to the second odor in the absence of shock. After 90 s of pure air, conditioning performance was tested for 120 s by exposing one side of the arena to the shock-associated odor and the other side to the neutral odor. (D) Complete conditioning protocol.

**Supplementary Figure 4.**
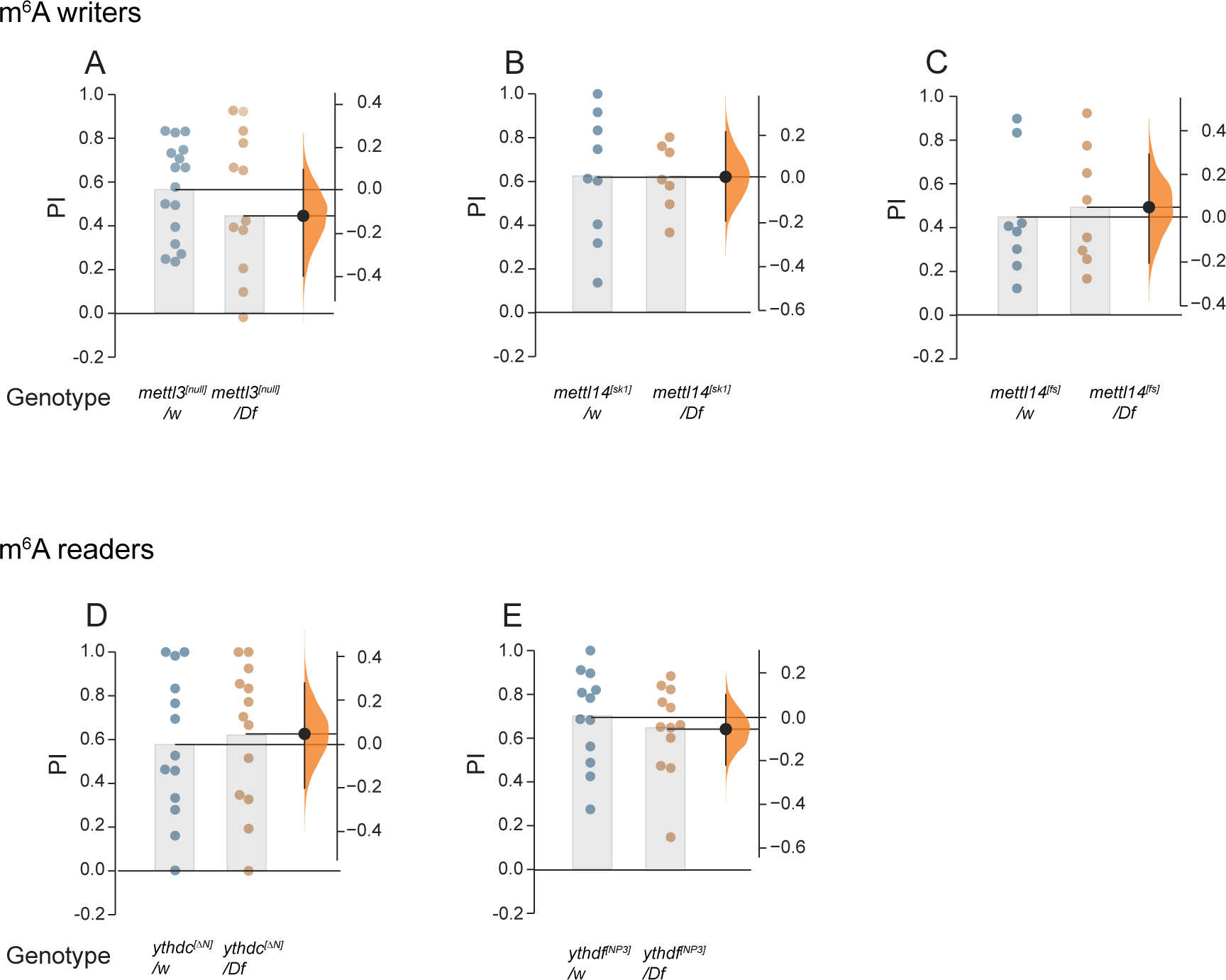
Minimal impairment of STM in 10 day old m^6^A pathway mutants (A) We observed a slight reduction in short term learning and memory (STM) in the odor avoidance paradigm in hemizygous *mettl3* mutants compared to heterozygous controls. *mettl3*^[null]^/Df vs *mettl3*^[null]^/w = -0.12[95CI -0.4, +0.1] *p* = 0.32. (B-E) No changes in STM performance were observed in other m6A pathway mutants. *mettl14*^[sk1]^/Df vs *mettl14*^[sk1]^/w = 0[95CI -0.2, +0.21] *p* = 0.9881. *mettl14*^[fs]^/Df vs *mettl14*^[fs]^/w = 0.04[95CI -0.21, +0.29] *p* = 0.7494. *ythdc*^[ΔN]^/Df vs *ythdc*^[ΔN]^/w = 0.05[95CI -0.2, +0.28], *p* = 0.706. *ythdf*^[NP3]^/Df vs ythdf^[NP3]^/w = -0.06[95CI -0.25, +0.18], *p* = 0.5961. N for each condition = >130 flies.

**Supplementary Figure 5.**
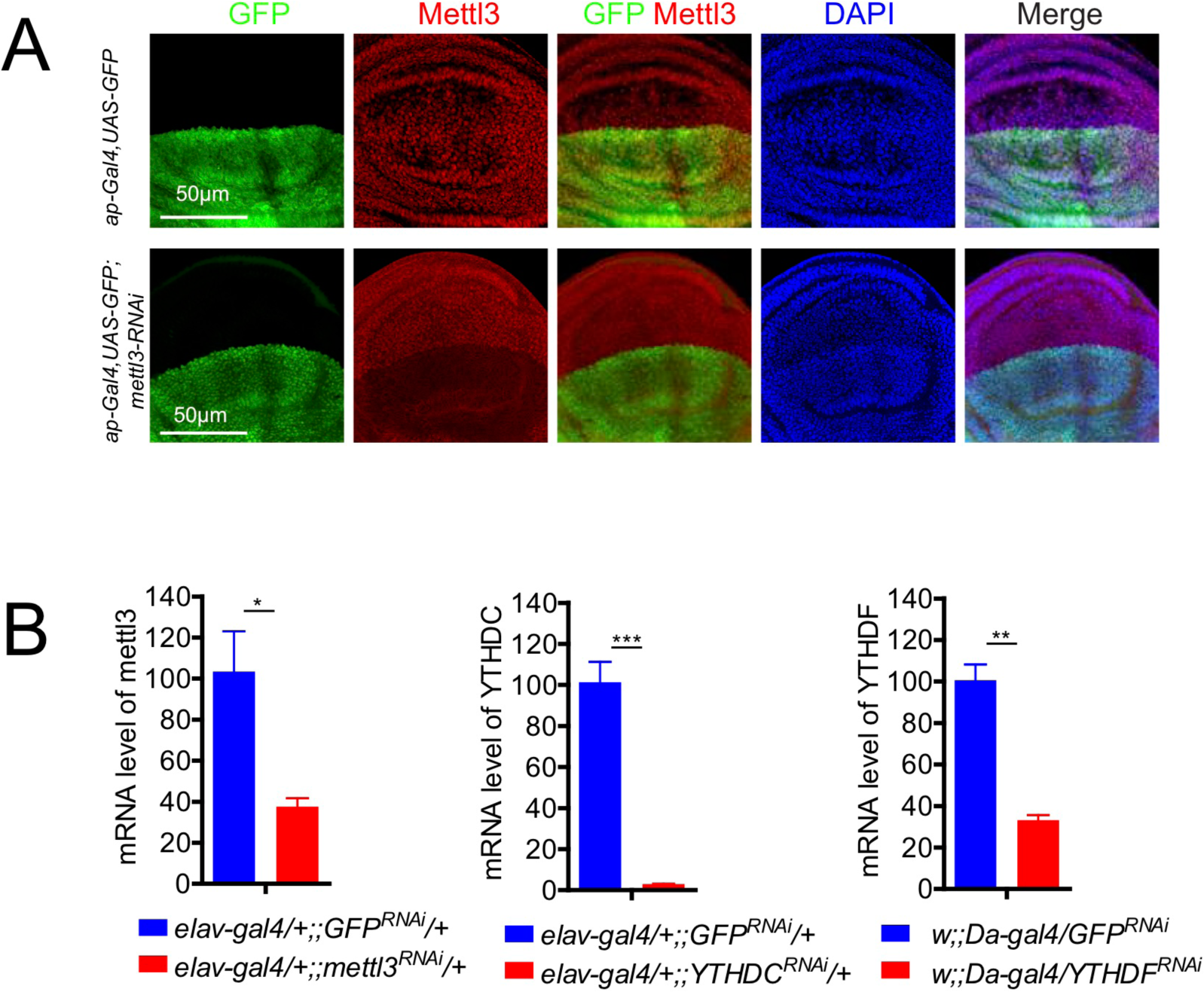
Validation of RNAi transgenes against m^6^A factors. (A) Wing imaginal disc pouch regions carrying *ap-Gal4>UAS-GFP* transgenes, stained for GFP (green) to mark the Gal4/knockdown territory, Mettl3 (red), and DAPI (blue). (A) Control disc shows relatively uniform nuclear Mettl3 signals. *ap-Gal4>UAS-Mettl3[RNAi]* disc shows specific loss of Mettl3 within the GFP+ dorsal compartment. (B) Validation of *UAS-mettl3[RNAi]*, *UAS-YTHDC[RNAi]*, and *UAS-YTHDF[RNAi]* transgene showing knockdown of cognate targets by qPCR.

**Supplementary Figure 6.**
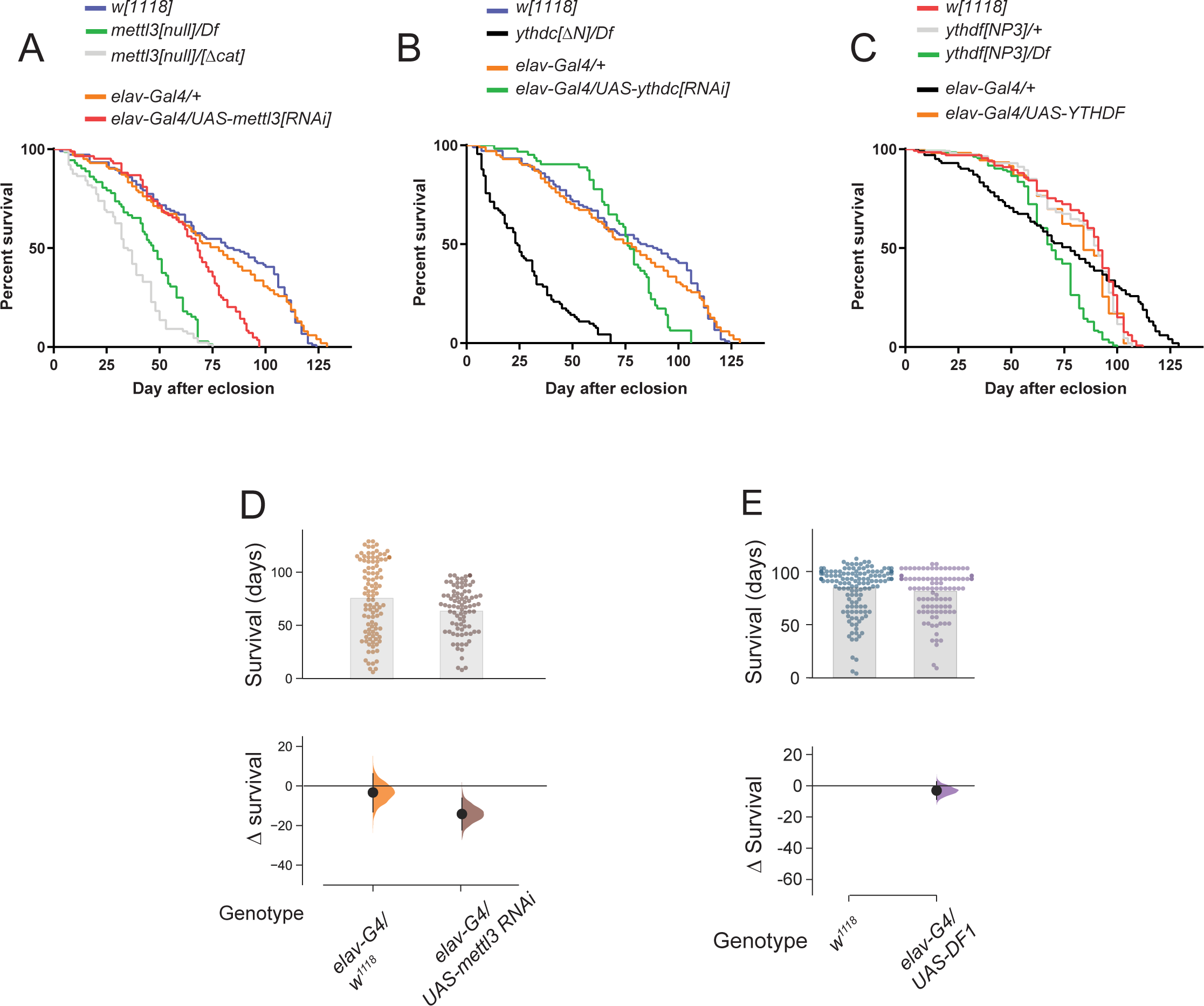
Lifespan measurements of m^6^A pathway manipulations. (A-C) Kaplan-Meier survival curves of adult *Drosophila* lifespan. These data correspond to scatter plots of adult survival shown in main Figure 3A-C. (A) Loss of *mettl3* reduced average *Drosophila* survival by >34 days compared to heterozygous control or wild type *w[1118]*. Neuronal-specific depletion of mettl3 led a moderate reduction of lifespan. (B) Loss of *ythdc* reduced *Drosophila* survival by about 50 days while *YTHDC* overexpression did not affect *Drosophila* lifespan. Neuronal-specific depletion of ythdc moderately reduce lifespan. (C) Loss of *ythdf* reduced *Drosophila* survival by about 14 days in the hemizygous background compared to heterozygotes and control wild type. All survival experiments were performed at 23°C with at least N=90 flies per condition. (D) Neuronal depletion of *mettl3* using *elav-Gal4* and *UAS-mettl3-RNAi* led to a moderate reduction in lifespan compared to the *elav-Gal4* driver alone. (E) Neuronal overexpression of YTHDF1 did not affect lifespan. In (D-E), the top plots show survival of each individual fly (dots). The height of the bar shows the average survival of the respective population. Bottom plot shows the delta survival of the respective genotype as compared to the control stocks (*elav-Gal4* or *w[1118]*).

**Supplementary Figure 7.**
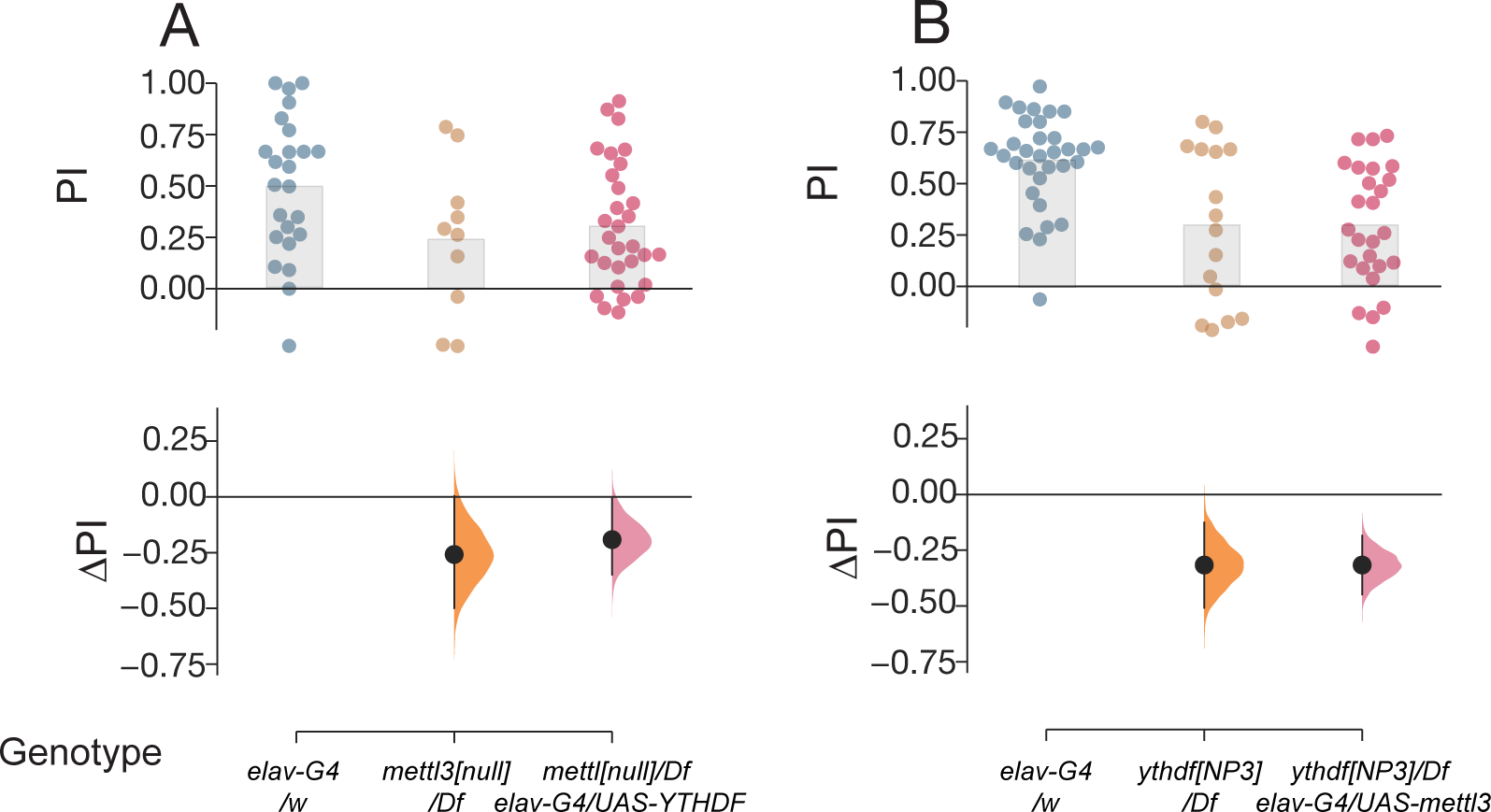
No cross-rescue capacity of *mettl3* and *YTHDF* in STM. (A) Pan-neuronal expression of *YTHDF* using elav-Gal4 (elav-G4) did not improve STM impairment in *mettl3* mutants. *mettl3*^[null]^/*Df* vs *elav-G4/w* = -0.26[95CI -0.5, 0], *p* = 0.0854. *mettl3*^[null]^*/Df, elav-G4/UAS-YTHDF* vs *elav-G4/w* = -0.19[95CI -0.35, 0], *p* = 0.0302 B: (B) Pan-neuronal expression of *Mettl3* in *ythdf* mutants did not rescue their STM impairment. *ythdf*^[NP3]^/*Df* vs *elav-G4/w* = -0.32[95CI -0.51, -0.13], *p* = 0.0105. *ythdf*^[NP3]^*/Df, elav-G4/UAS-mettl3* vs *elav-G4/w* = -0.32[95CI -0.45, -0.18], *p* = 5*10^-5^. N for each condition > 120 flies.

**Supplementary Figure 8.**
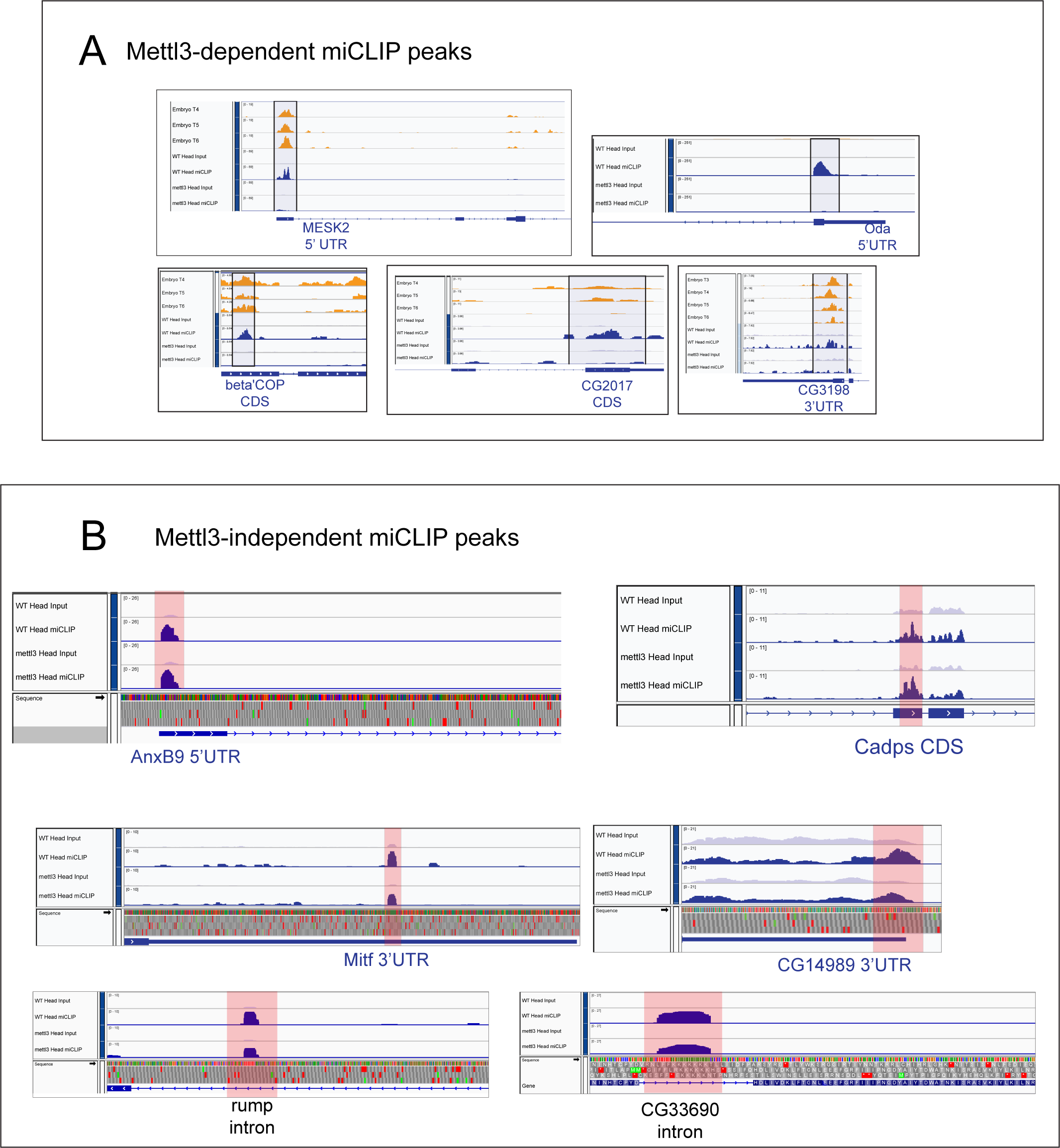
Examples of Mettl3-dependent and -independent m^6^A peaks. (A) IGV screenshots of several targets displaying Mettl3-dependent miCLIP peaks (highlighted in blue) in various genomic locations (5’UTR, CDS and 3’ UTR). Embryo miCLIP data are from Kan et al (2017) and have substantial concordance with our new data at Mettl3-dependent loci even though the tissue types are distinct. (B) IGV screenshots of Mettl3-independent miCLIP peaks (highlighted in red) in various genomic locations (5’UTR, CDS, introns and 3’ UTR)

**Supplementary Figure 9.**
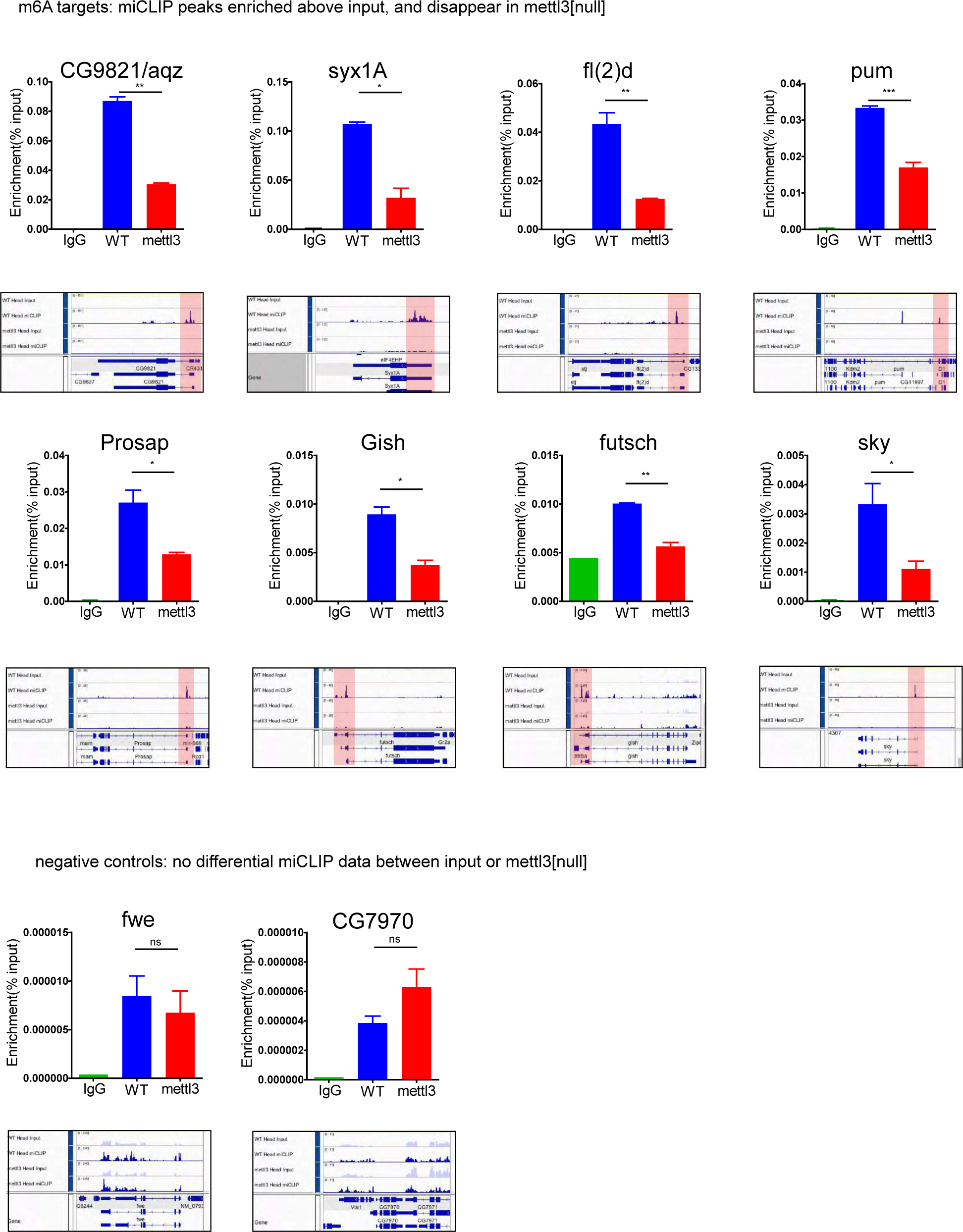
m^6^A target transcripts selected for validation. Shown are IGV screenshots showing Mettl3-dependent miCLIP peaks (highlighted in pink boxes) at genes selected for m^6^A-IP validation followed by rt-qPCR; enrichments are shown in IgG control, wild type and *mettl3[null]* flies. Negative control genes lacking m^6^A peaks are shown at bottom.

**Supplementary Figure 10.**
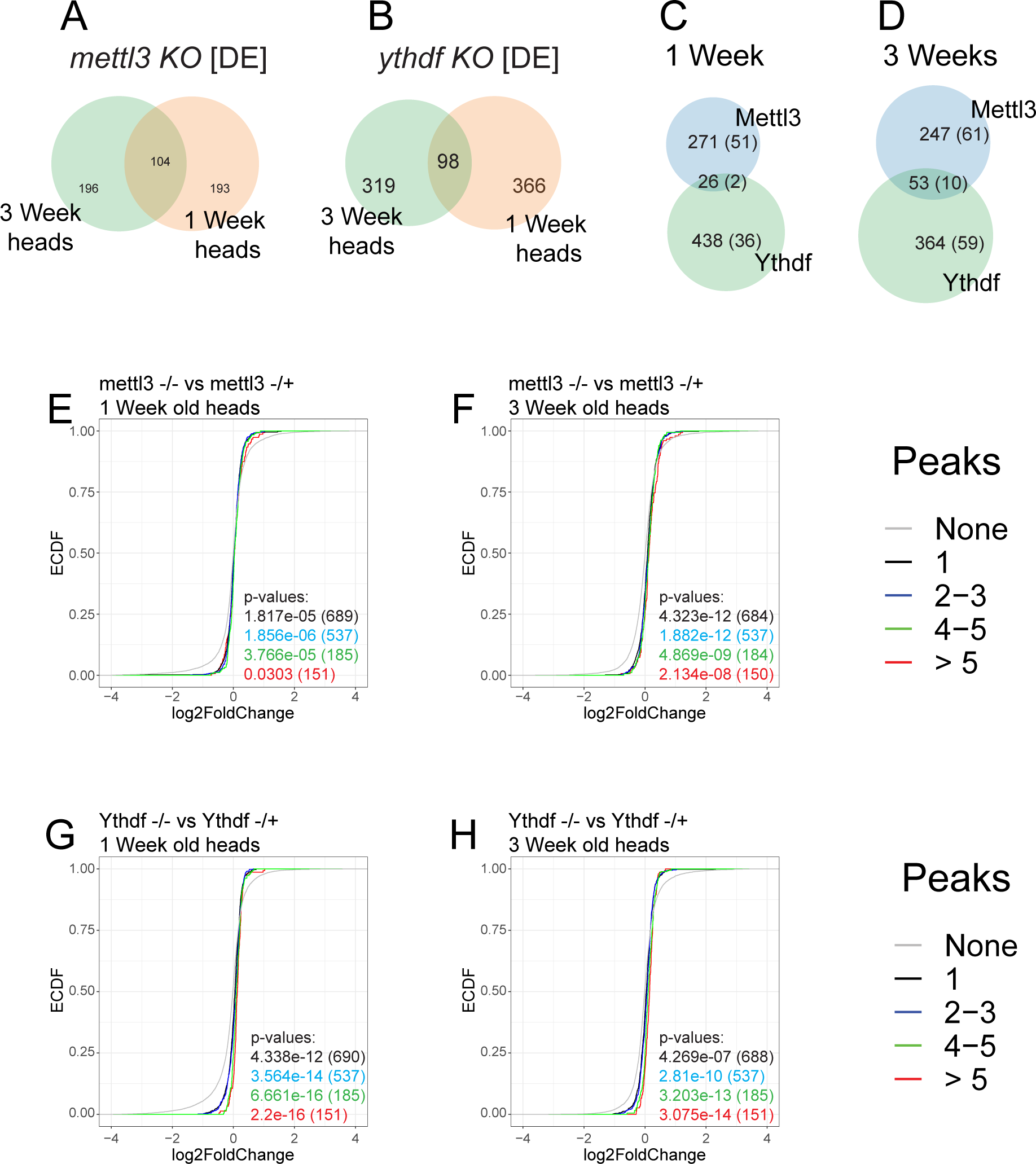
m^6^A target genes are not differentially expressed (DE) under writer or reader knockdown in the CNS. (A-B) Overlap in consistently differentially expressed genes at 1- and 3-week time points in writer (mettl3 - A) or reader (ythdf - B) knockout heads. (C-D) Overlap between consistently DE genes in writer (mettl3) and reader (ythdf) knockout heads at 1-week (C) and 3-week (D) timepoints. Numbers in parentheses are m6A target genes. (E-H) No directional change observed in m^6^A target genes in writer (E and F) or reader (G and H) mutants at 1-week (E and G) or 3-week (F and H) timepoints. Cumulative distribution plots were generated by grouping m^6^A target genes based on numbers of peaks per gene. Comparisons are listed above each plot. A bootstrap method generated the background distribution (None) using genes that lacked m^6^A peaks. To generate p-values, two-sided Kolmogorov-Smirnov (KS) tests were performed comparing the background distribution and each group of m6A target genes. Number of targets are included in parentheses.

**Supplementary Figure 11.**
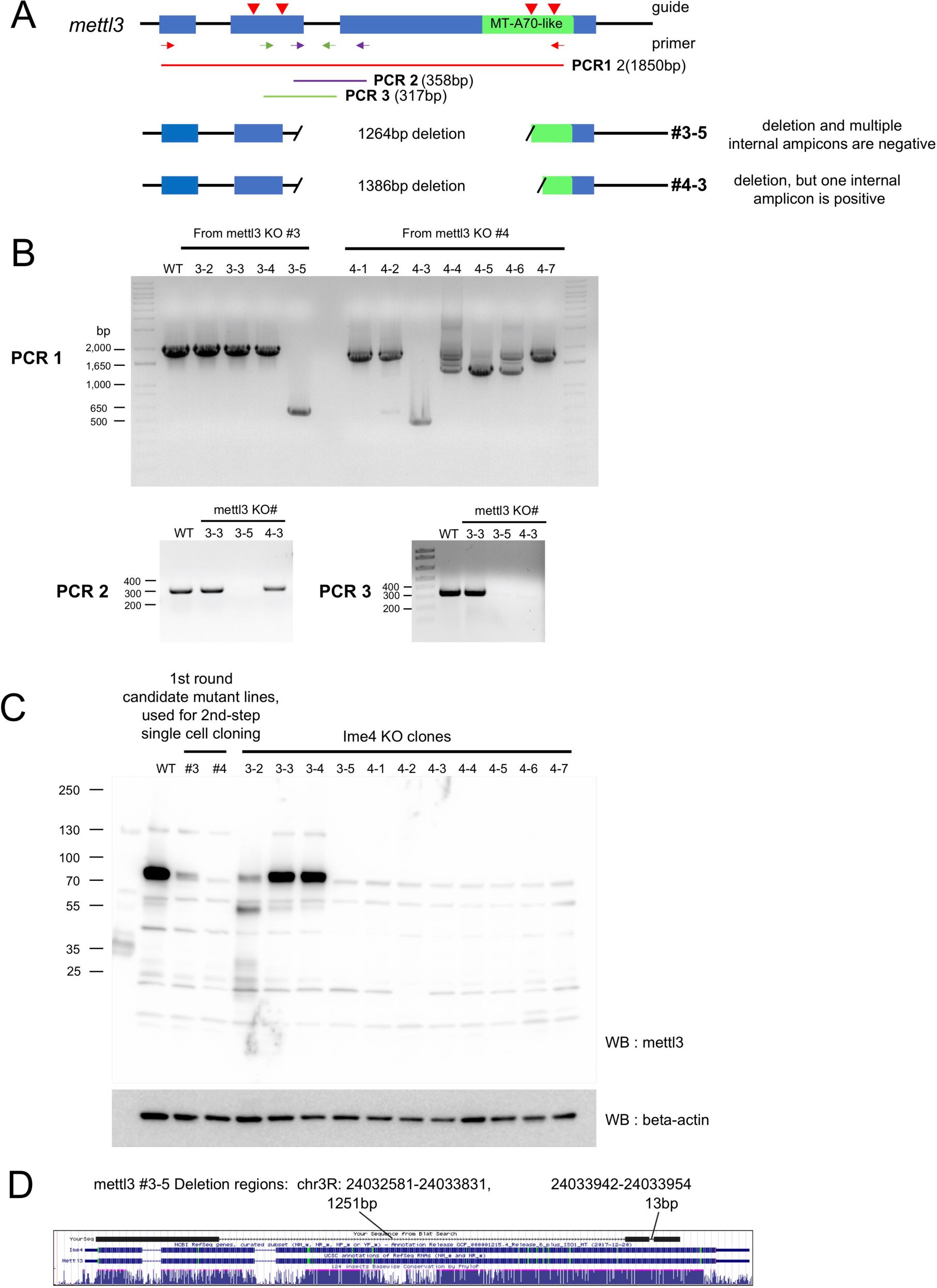
Generation of *mettl3-KO* cell lines. (A) Guide RNAs and multiple amplicons used to genotype *mettl3-KO* cell are shown. Two lines with large deletions were isolated, but one of these (#4-3) retained amplification of an internal amplicon even though it was apparently protein null. (B) PCR screening of *mettl3-KO* clones. Because S2 cells are difficult to grow clonally, we initially grew limiting dilutions of puro-Cas9/sgRNA cells that survived drug selection with candidate deletions, and then reselected clonal lines with deletions, and further characterized lines #3-5 and #4-3. (C) Western blotting of Mettl3 protein confirms *mettl3-KO* clones. (D) Deletion regions of *mettl3-KO* #3-5 determined from sequencing; this clone was used for mass spec validation (see main Figure 5D).

## Supplementary Tables

Supplementary Table 1. Proteomics Data

The proteomics data file for Figure 1C (searched against *Drosophila melanogaster* database UniProt (https://www.uniprot.org/).

Supplementary Table 2. Overview of miCLIP and accompanying datasets

Table lists miCLIP and input libraries reported in this study. Counts of pre- and post-map read processing, mutation calling and annotation of CIMs are listed as described in Grozhik et al (Grozhik et al., 2017).

Supplementary Table 3. Mettl3-dependent miCLIP peaks

Table contains a list of Mettl3-dependent, split peaks. Metadata include enrichment scores, gene segment annotation.

Supplementary Table 4. CIMs calls from miCLIP head libraries.

Table contains the locations of putative single nucleotide m^6^A sites using the miCLIP CIMs analysis pipeline detailed in Grozhik et al (Grozhik et al., 2017). Score metadata reflects m/k ratios described in the methods section.

Supplementary Table 5. Differential gene expression analysis

Table lists differentially expressed genes in 1 and 3 week *mettl3-/-* and *ythdf-/-* fly head libraries. Sheet names detail comparisons and metadata includes fold change and measurements used to determine statistical significance.

Supplementary Table 6. Oligonucleotide sequences.

This file contains the sequences of qPCR primers, genotyping primers, guide RNA, and oligos for making constructs used in the study.

